# Combining Ancient DNA and Radiocarbon Dating Data to Increase Chronological Precision

**DOI:** 10.1101/2020.09.18.300087

**Authors:** Jakob W. Sedig, Iñigo Olade, Nick Patterson, David Reich

## Abstract

This paper examines how ancient DNA data can enhance radiocarbon dating. Because there is a limit to the number of years that can separate the dates of death of related individuals, the ability to identify first-, second-, and third-degree relatives through aDNA analysis can serve as a constraint on radiocarbon date range estimates. To determine the number of years that can separate related individuals, we modeled maximums derived from biological extremes of human reproduction and death ages and compiled data from historic and genealogical death records. We used these estimates to evaluate the date ranges of a global dataset of individuals that have been radiocarbon dated and for which ancient DNA analysis identified at least one relative. We found that many of these individuals could have their date ranges reduced by building in date of death separation constraints. We examined possible reasons for date discrepancies of related individuals, such as dating of different skeletal elements or wiggles in the radiocarbon curve. Our research demonstrates that when combined, radiocarbon dating and ancient DNA analysis can provide a refined and richer view of the past.

## 1. Introduction

This article examines how aDNA data can be used innovatively to help with a central aspect of archaeological research—chronology. Ancient DNA (aDNA) data are revolutionizing the field of archaeology. Within the last decade alone, aDNA analyses have discovered new hominins (Reich et al., 2010), elucidated the spread of farming through Europe (Lazaridis et al., 2016; Mathieson et al., 2015), shed light on the peopling of the Americas and Oceania (Lipson et al., 2018; Moreno-Mayar et al., 2018; Posth et al., 2018; Rasmussen et al., 2014; Skoglund et al., 2016), and more. While aDNA has helped provide insight on long-standing archaeological questions, exponentially increasing aDNA data has created unique opportunities for the examination of finer-grained issues, and even archaeological methods.

The basis of the work presented here is tied to the fact that there is a maximum number of years that can separate the dates of death (DOD) for two or more genetically related individuals. For example, it is exceedingly rare for a mother to die 100 years before her daughter, particularly in pre-modern societies. Thus, if two or more individuals are identified as biological relatives through aDNA analysis and those individuals are radiocarbon dated, their relatedness can be used as a prior or constraint when analyzing their overlapping radiocarbon date ranges. Using these constraints, we examine how the identification of genetic relatives can help identify errors and outliers in radiocarbon dating, how biological relatedness can be used to constrain overlapping radiocarbon date ranges and increase dating precision, and how application of the methods to a large database of published ancient DNA data (https://reich.hms.harvard.edu/downloadable-genotypes-present-day-and-ancient-dna-data-compiled-published-papers) can reveal potential larger issues in the radiocarbon record at particular times and places.

## 2. Materials and Methods

### 2.1 Identification of genetic relatives with ancient DNA

Identification of genetic relatives has become standard practice in ancient DNA analysis. Typically, individuals which are screened and produce working genomic data are compared against each other and previously analyzed individuals from similar geographic regions and time periods to identify unique genetic relationships. For each pair of individuals in this study, we computed the mean mismatch rate using all the autosomal SNPs with at least one sequencing read for both individuals in the comparison (this procedure to identify genetic relatives is described in Kennett et al. (2017:156) and van de Loosdrecht et al. (2018:15), and is similar to that in Kuhn et al. (2018:157)). In the cases with more than one sequencing read at a particular SNP for a given individual, we randomly sample one for analysis. We then estimate relatedness coefficients as in Kennett et al (2017:156): r = 1–((x-b)/b) with x being the mismatch rate and b the base mismatch rate expected for two genetically identical individuals from that populations, which we estimate by computing intra-individual mismatch-rates. We also compute 95% confidence intervals using block jackknife standard errors (Olalde et al., 2019:S61). While such analysis can detect relationships up to the 5th degree, we limit relationships here to 3rd degree maximum, as DOD date separations become too great to be of use with decreasing genetic relatedness (e.g. great-grandparents and grandchildren).

### 2.2 Genetic relatives and DOD separation maximums

Below, two approaches—biological maximums and genealogically and historically derived estimates—are examined for determining the DOD separation of genetically related individuals. The biological maximums serve as theoretical extremes that, while biologically possible, are very rare and unlikely to occur, especially in pre-industrial archaeological cultures. Genealogically and historically (GH) derived DOD separations were created through the examination of genealogical records and historic data and reflect more realistic estimates of the number of years between the death of two related individuals.

### 2.3 Biological Maximum DOD Estimates

Biological maximum estimates use extremes of human reproduction and lifespan to produce maximum DOD separation estimates. Figures 1 and 2 (also see SM 1) are diagrams of how these estimates were modeled. The start of these models is set at 0 CE. At this point, a couple consisting of a 15-year-old male and female parent a male child. This child dies at birth, but both parents live to be 100 years old. Thus, the DOD separation between the child and parents would be 85 years. If instead the mother died during childbirth, but the child lived to 100 years old, the maximum DOD separation between parents-offspring would be 100 years. Siblings have an even greater potential maximum DOD separation, as Figure 2 demonstrates. In this model, the 15-year-old couple has a male child that dies at 0 CE. That same couple has another child 30 years later (when they are 45 years old); that second child then dies 100 years later. So, the maximum separation between the siblings is 130 years.

**Figure 1.**
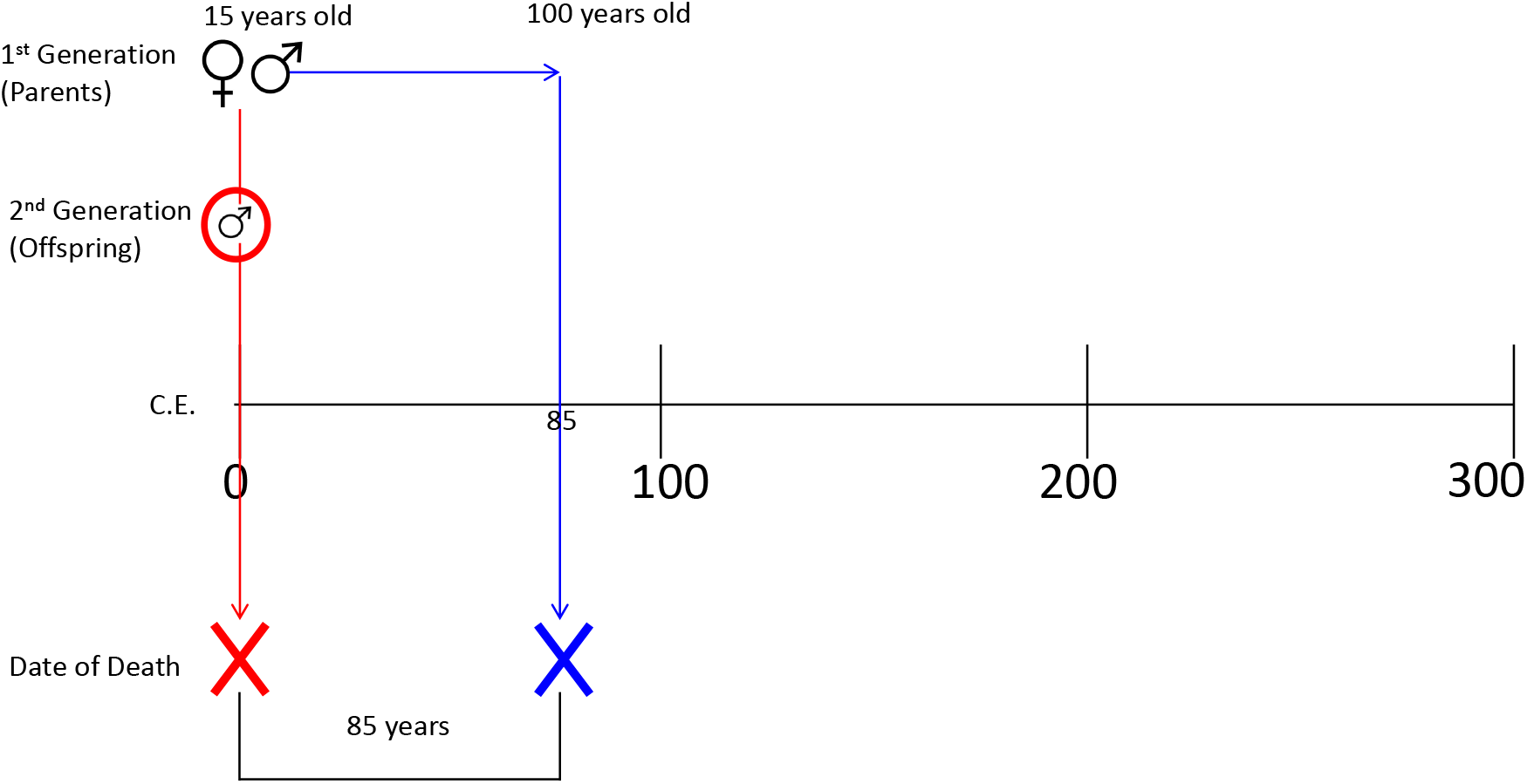
Model of biological maximum date of death separation for parents-offspring.

**Figure 2.**
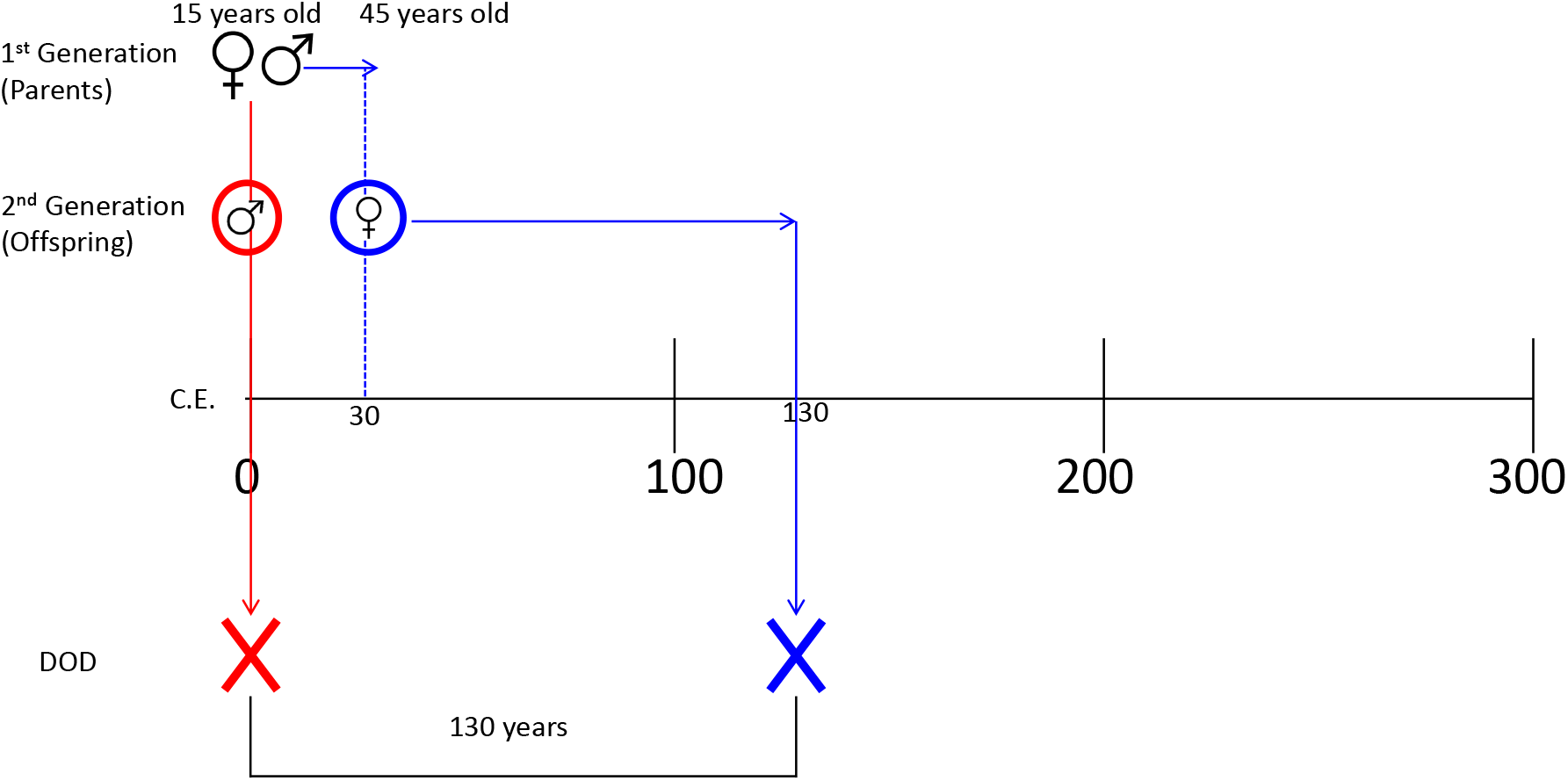
Model of biological maximum date of death separation for siblings.

Using these parameters, a number of potential biological maximums were modeled for various degrees of genetic relatedness (Table 1; see SM 1 for diagrams of models). The biological DOD maximums presented above and in Table 1 are reliant on extremes—producing children at the biologically earliest and latest possible ages and living to extreme old age. While possible, these DOD separations are not realistic, and are largely ineffective as constraints on C14 date range distributions. Thus, to more effectively examine how date of death separations for related individuals can be applied to overlapping radiocarbon ranges, we also compiled birth and death data from historical and genealogical records.

**Table 1.**
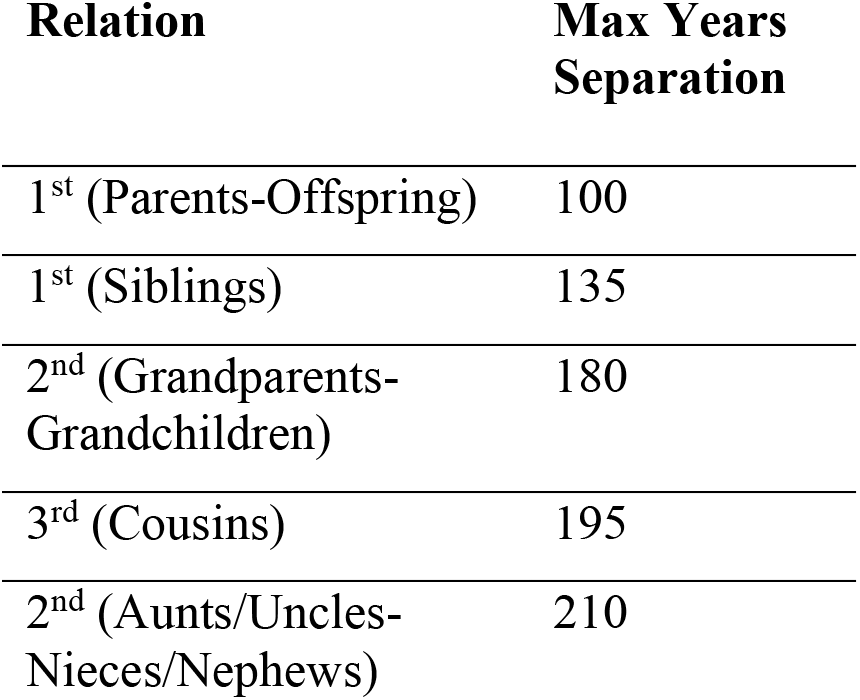
Theoretical DOD separation biological maximums.

### 2.4 Genealogically and Historically Derived DOD Estimates

We began compiling data on the date of death separations for related individuals by consulting the plethora of genealogical and historical data that are publicly available online. Many of these databases consist primarily of people of European ancestry who lived within the last two centuries. However, to create date of death estimates from heterogenous data, we sought non-European focused databases for relatives’ death dates. Data were gathered from historic Anglo cemeteries, and online databases of birth and death dates for Cherokee, Tlingit, and other Native American groups (SM 2). Data were sorted by categories of relatedness: parent-offspring, sibling, grandparent-grandchild, and other 2^nd^-3^rd^ degree (aunts-uncles/nieces-nephews and cousins).

The DOD separation for related individuals was compiled into a spreadsheet for each genealogical database (SM 2). DOD separations were calculated by identifying related individuals then subtracting the dates of death (i.e. if a mother and daughter were identified, and the mother died in 1800 CE and the daughter 1850 CE, the separation between the two entered in the database would be 50). For parent-child and grandparent-grandchild relationships the signed value of the DOD was recorded. As will be discussed later, knowing whether the child died before the parent (which would result in a negative value) is useful for building constraints of parent-child and grandparent-grandchild radiocarbon ranges. However, since in many instances aDNA cannot determine the relatedness direction of two individuals (e.g. which is the mother and which is the daughter) the absolute value of DOD separation of each relative pair was recorded for each relationship type and is primarily used for the analyses below.

A total of 5235 relative DOD separations were recorded: 800 parent-offspring, 813 sibling, 485 grandparent-grandchild, and 3137 other 2^nd^-3^rd^ degree. The means, medians, and standard deviations of the absolute value for each relationship type were then calculated; the results are provided in Table 2 (see also SM 2).

**Table 2.** Compiled genealogical and historical data for DOD absolute value separation

|  | **Parent-**<br>**Offspring** | **Sibling** | **Grandparent-**<br>**Grandchild** | **Other 2<sup>nd</sup>-3<sup>rd</sup>**<br>**relationships** |
| --- | --- | --- | --- | --- |
| Mean | 28.84 | 26.33 | 35.00 | 34.94 |
| Median | 26 | 20 | 39 | 30 |
| Standard Dev. | 18.94 | 22.41 | 31.93 | 25.6 |

The data in Table 2 demonstrate that the biologically maximum DOD separation estimates in Table 1 are truly extremes. The largest mean separation in the GH dataset was 35.00 years between grandparents-grandchildren. The single greatest DOD separation in all the data was 117 years between Cherokee 2^nd^/3^rd^ degree relatives—still 93 years short of the 2^nd^/3^rd^ degree maximum theoretical estimate (210 years). The mean GH DOD separation estimates for parents-offspring, siblings, and grandparents-grandchildren are 71.16, 108.67, and 145.00 less than the biological maximum separation estimates (Table 1), respectively.

The DOD separations above were produced by manual collection from online, publicly available resources. However, in a 2018 study Kaplanis and colleagues developed software and an analysis pipeline to examine genealogies of millions of individuals downloaded from the online genealogical database geni.com. Kaplanis et al. (2018) used this data to construct family trees (sometimes containing millions of individuals); the anonymized data from this study were made available to download (https://familinx.org/). Significantly, the data contained information on which individuals had parent-offspring relationships, and the death date for each individual. We therefore downloaded these data and found the DOD separation for over 8 million parents and offspring (SM 3). We removed pairs with data errors (for example a death date of 3500) and used the biological DOD separations defined above for parent-offspring as constraints (i.e. 85 years for children dying before parents and 100 years for parents dying before children). The mean absolute value DOD separation for these 8 million parent-offspring pairs was 31.43 years, slightly higher than the mean value we manually collected (28.84; SM 2); however, this should be expected as the geni.com data is heavily weighted toward modern, European individuals who likely had longer life spans. Overall, the similarity between the Kaplanis et al. 2018 data and the genealogical and historical data we manually curated demonstrates that the DOD separations we obtained represent more realistic DOD separations for related individuals than the biological maximums.

Although the DOD separation estimates derived from GH data are more reflective of separations between genetic relatives than the biologically possible maximums, the GH data presented here should be viewed only as rough estimates. More precise estimates could be tailored for particular types of social organization, such as hunter-gatherers, pastoralists, agriculturalists, city-dwellers, nomads, etc. However, should researchers wish to create new models, the GH estimates above likely will not be exceeded, as many of the separations were derived from individuals who lived after the industrial revolution and likely had longer lifespans than ancient individuals.

## 3. Application and analysis

### 3.1 Applications of relatedness data to radiocarbon dated individuals

We examined the ancient DNA database of published individuals from geographic locales across the globe spanning more than 30,000 years (although there is bias towards the last 10000 years in western Eurasia; see Marciniak and Perry, 2017; Reich, 2018) to test how DOD separation estimates can be applied to related individuals and examine if any new insights can be revealed. As of May 2020, 3,965 published individuals were in the database, with 1,127 ancient individuals having at least one identified relative. Of those, 190 pairs (231 unique individuals, SM 4) had both individuals C14 dated (all dates generated with AMS and calibrated two-sigma), allowing for analysis of DOD separations and constraints.

### 3.2 Outlier identification

The most basic example of how genetic relatedness can help refine radiocarbon dating is through the identification of anomalies. Archaeologists have long recognized that C14 sample contamination can occur and that other issues, such as the marine reservoir effect, can cause dates to be skewed (Taylor and Bar-Yosef, 2014). Genetic relatedness is a new independent measurement that can be used to test the validity of radiocarbon date ranges, particularly for samples that might not be obvious outliers. For example, if five skeletons from the same stratigraphic layer in a cemetery were dated, and four of those individuals had calibrated ranges of approximately CE 1-500, while one had a calibrated date range of approximately 2500-2000 BCE, that one sample would seem suspicious and would likely be redated (Figure 3). However, if that outlier instead had a range of approximately CE 600-1000, the skeleton might not be redated, as it is relatively close to the range of the other four skeletons (Figure 3). But, if it was determined that the outlier was actually the father of skeleton 3, then it would be highly suspicious that skeleton 5 could be older than skeleton 3, as the DOD separation between father and offspring cannot biologically be more than 100 years, and more realistically is around 29 years from GH DOD estimates (Table 1). Such an example was discovered in the database.

**Figure 3.**
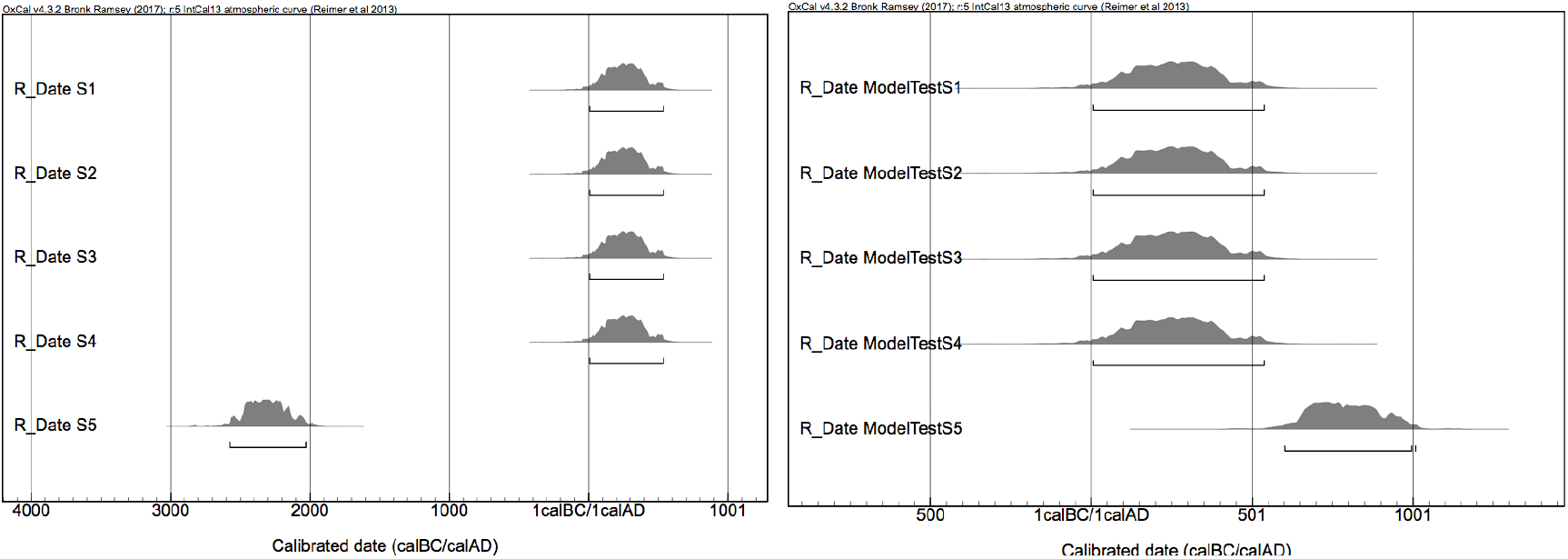
Examples of clear outliers in radiocarbon dating (L) and an individual that is an outlier but does have overlap with other dates (R).

Two individuals, I2457 and I2600 (Olalde et al., 2018), were excavated from two sites, Amesbury Down and Porton Down, Britain, separated by approximately 5km. The samples had been previously radiocarbon dated; prior to aDNA analysis and the dates did not seem suspect (I2457= 3890+30; 2480-2280 calBCE, SUERC-36210; I2600=3646+27; 2140-1940 calBCE, SUERC-43374; Figure 4). Ancient DNA analysis of the samples revealed that I2600 was the daughter of I2457, but there was thus no overlap in the calibrated distributions of the father-daughter pair. The minimum DOD separation between the father and daughter was 140 years, which exceeds even maximum biological estimates. Individual I2457 (the father) was therefore redated and the new date (3717+28; 2200-2031 calBCE; SUERC-69975) fit within the expected DOD spread (Figure 5).

**Figure 4.**
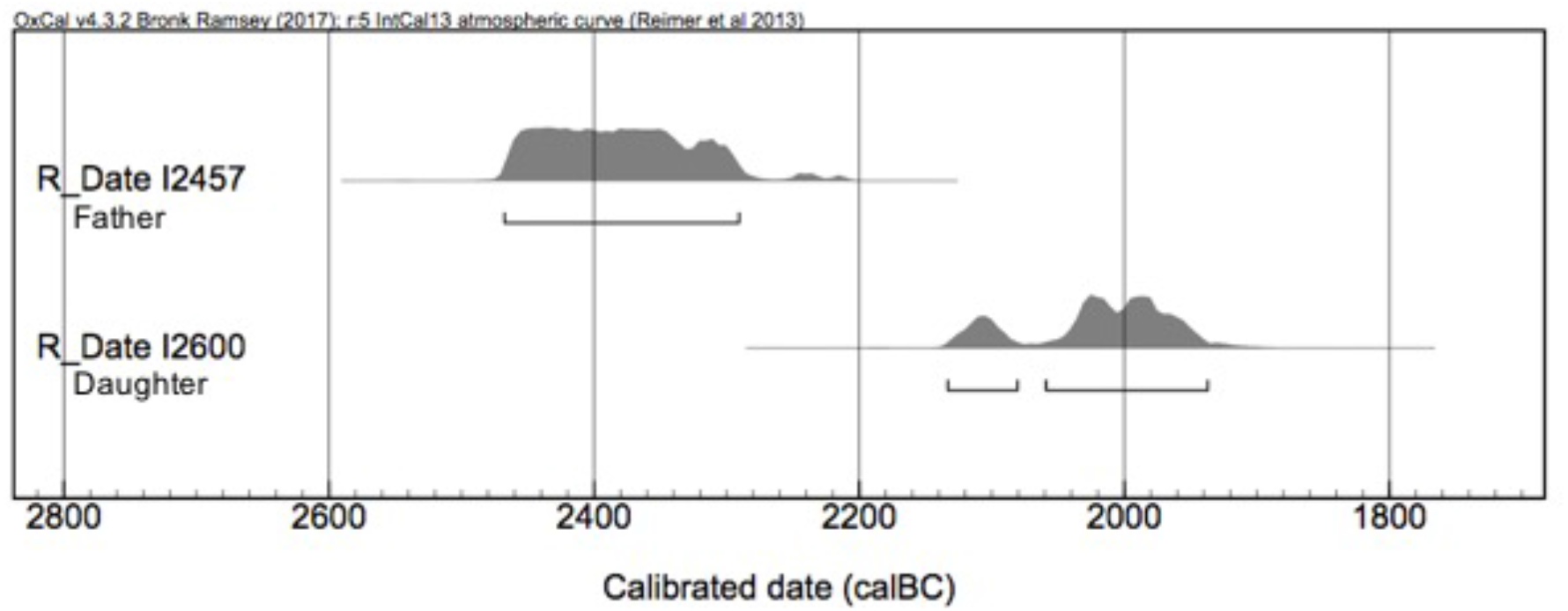
Original AMS dates for I2457 and I1600.

**Figure 5.**
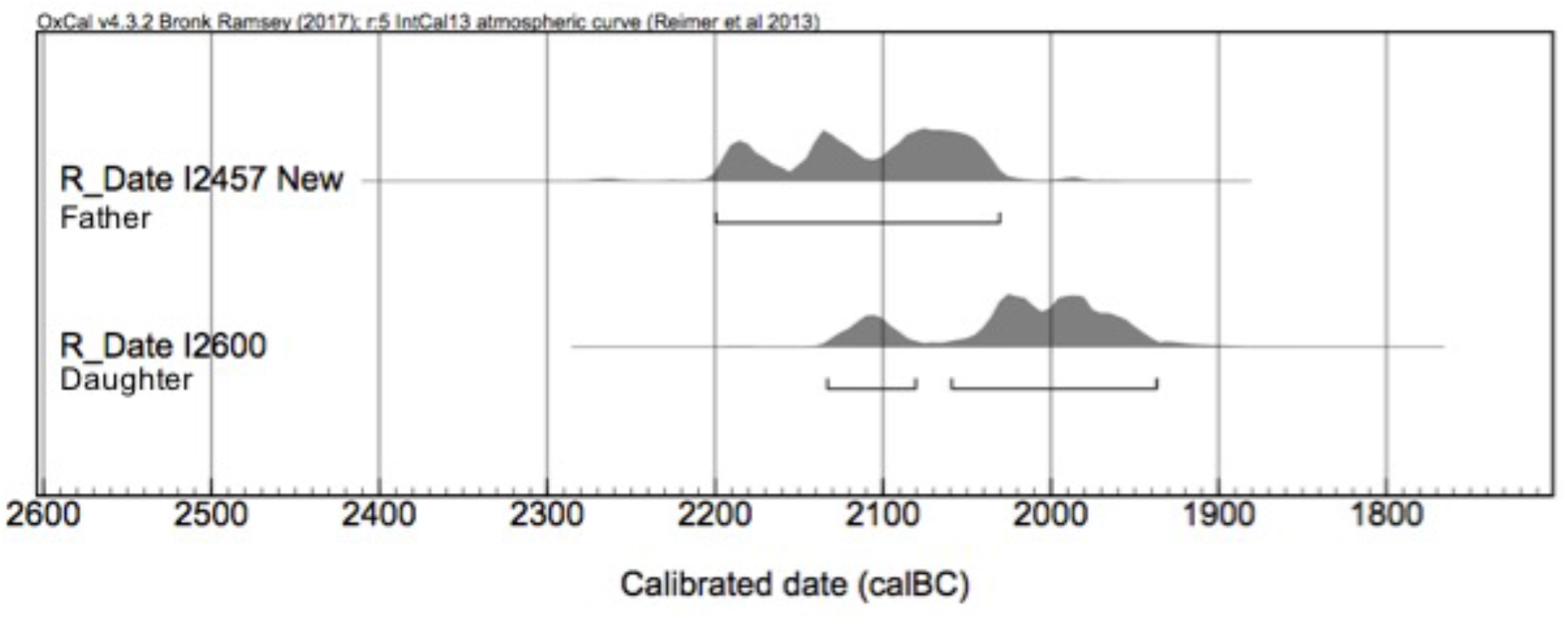
New AMS date for I2457.

### 3.3 Range tightening of radiocarbon date distributions

Along with detecting outliers, we examined the potential of using DOD separations to refine calibrated date ranges in instances where two (or more) related individuals with overlapping C14 date probability distribution ranges are identified. Using biological maximums as an example, consider a father whose AMS range is 1-500 calCE and a daughter whose range is 400-1000 calCE. Since the father cannot have died more than 100 years before or after the daughter, the father’s range can be constrained to approximately 300-500 CE; since the daughter cannot have died more than 100 years after the father, the daughter’s range can be constrained to 400-600 CE.

While informative for some related pairs, the maximum biological separation estimate is often too large and not applicable to most related and dated individuals in the dataset. Individuals I2457 and I2600 serve as examples of how GH DOD separations can be used as constraints for the date ranges of related individuals. Using the new date for I2457 and building in the 29-year parent-offspring GH DOD constraint allows the individual and combined date ranges to be reduced (Figure 6). With this estimate, the tail ends of the calibrated distributions for I2457 and I2600 should not be separated by more than 29 years. In other words, since I2457 likely died an average of 29 years before I2600, adding 29 years to the left end of the two-sigma calibrated date range for I2600, 2134 BCE, creates a constraint for the earliest date of I2457 at 2163 BCE. On the other end of the distributions, since I2600 likely did not die more than 29 years after I2457, adding 29 years to the latest dates of the 2-sigma calibrated range of I2457, 2031 BCE, creates a constraint for the latest date of I2600 at 2002 BCE (Figure 6).

**Figure 6.**
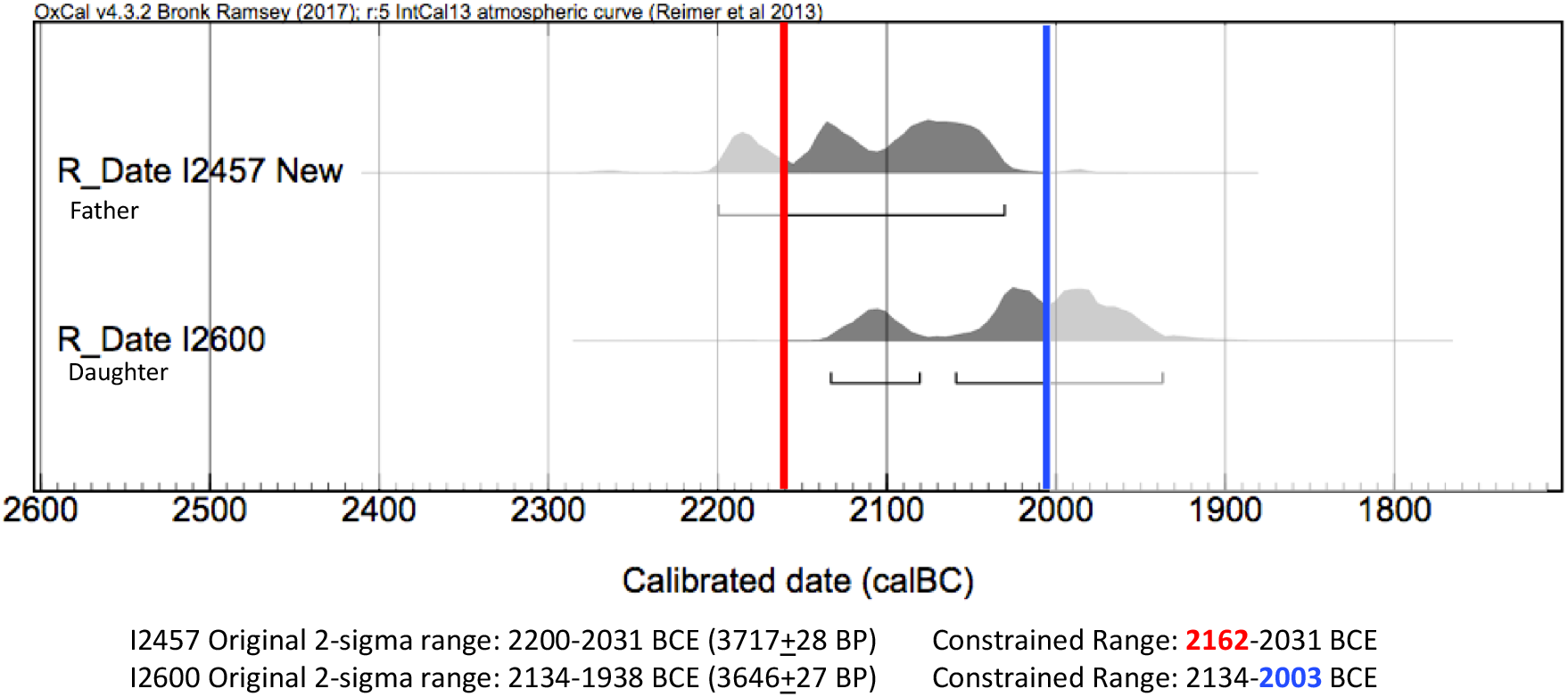
AMS ranges for I2457 and I2600 with relative constraints added.

Not all date ranges can be constrained as significantly as with these relatives, as demonstrated by individuals I1054 and I1053 (Narasimhan et al., 2019), two siblings from the Russian Sintashta archaeological culture. Their date ranges overlap almost entirely (Figure 7). Thus, using the 26-year GH DOD separation estimate for siblings, only a minor reduction can be made for their individual and combined 2-sigma date ranges (a change in only two years).

**Figure 7.**
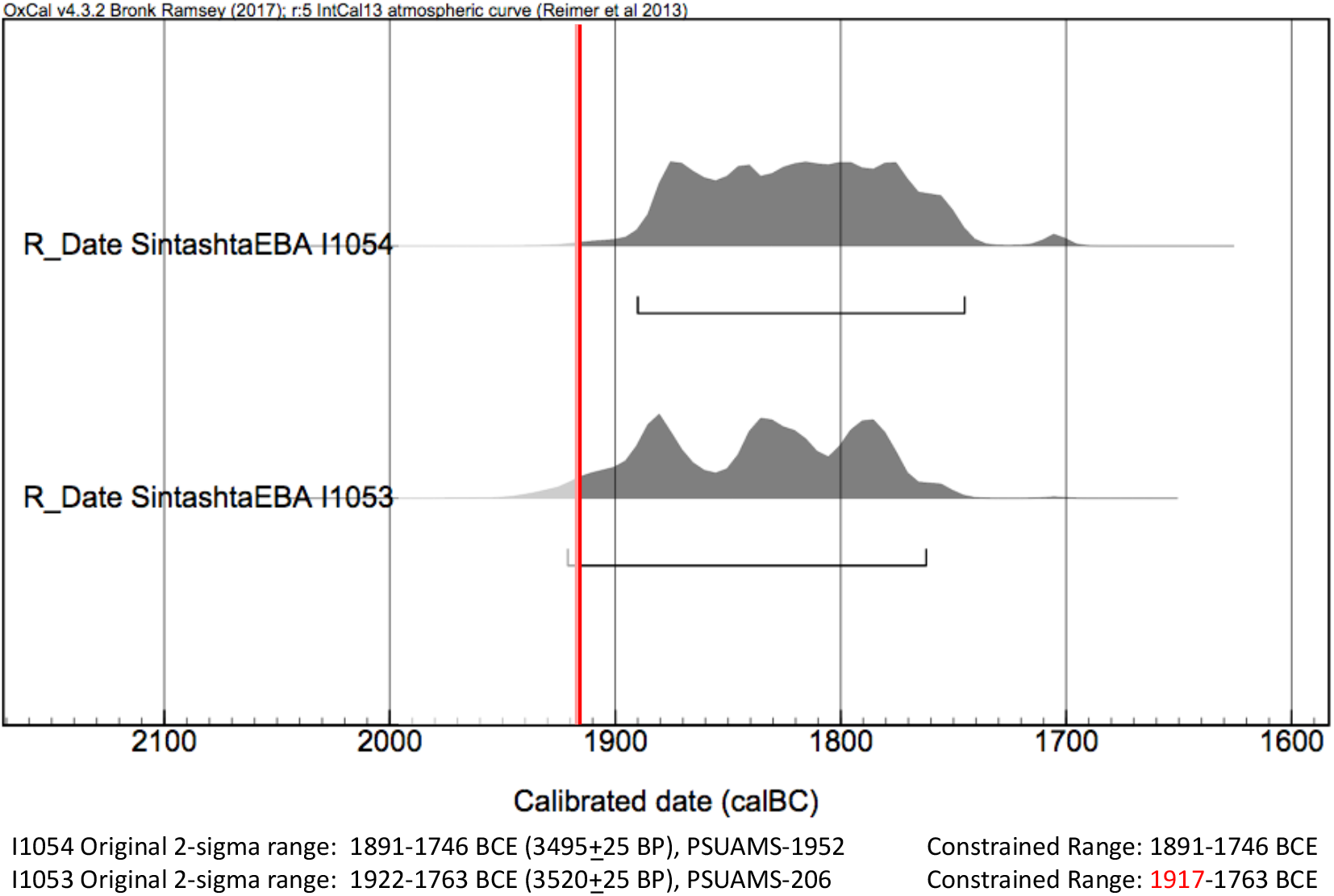
AMS ranges for I1054 and I1053 with relative constraints added.

### 3.4. Constraints applied to database

We applied the date range distribution tail trimming approach outlined above to the 190 dated relative pairs in the database (we used separations of 29 years for parent-offspring, 26 for siblings, and 35 years for 2nd-3rd degree of unknown specificity as constraints). We focus below on GH DOD derived constraints, as the biologically maximal DOD constraints often exceeded the overlap of the related pairs’ C14 distributions. As mentioned above, since the type and directionality of relationships often cannot be precisely determined through aDNA analysis, we used the largest mean absolute values for DOD separations derived from GH data. In other words, if a pair could only be distinguished as 1st degree relatives (either parent-offspring or siblings), the largest mean GH DOD separation for first degree relatives was used, which is 29 years (for parent-offspring, not 26 years for siblings).

After applying GH DOD constraints to the dataset, we removed 21 pairs because their 2-sigma calibrated date ranges did not overlap and exceeded the GH DOD estimates, suggesting dating error/a need for redating (uncorrected marine reservoir effect, sample contamination, etc). This left a total of 169 pairs and 219 unique individuals (SM 4). Applying the GH constraints, we were able to reduce the 2-sigma calibrated ranges of 132 individuals, with a mean reduction of 54.47 years (Table 3 and SM 4.2); 77 individuals had even more of a reduction than this. Figure 8 is a graph of the difference between the original 2-sigma calibrated range and the GH constrained range for each individual.

**Table 3.** Date ranges for all individuals in relative pairs.

|  | **Original**<br>**Range<sup>1</sup>** | **GH**<br>**Range<sup>2</sup>** | **(O)-**<br>**(GH)**<br>**Change<sup>3</sup>** |
| --- | --- | --- | --- |
| Mean | 203.44 | 149.00 | 54.47 |
| Standard<br>Deviation | 95.89 | 86.60 | 75.82 |

<sup>1</sup>Original range is the 2-sigma calibrated range for the individual. <sup>2</sup>GH range builds in the constraints derived from ethnographic/historic records for each individual (Table 2).
<sup>3</sup>(O)-(GH) Change is the original range minus the constrained biological range for each individual, which reveals how many years of the original range are removed from the tail ends of the 2-sigma distributions when GH estimate constraints are applied.

**Figure 8.**
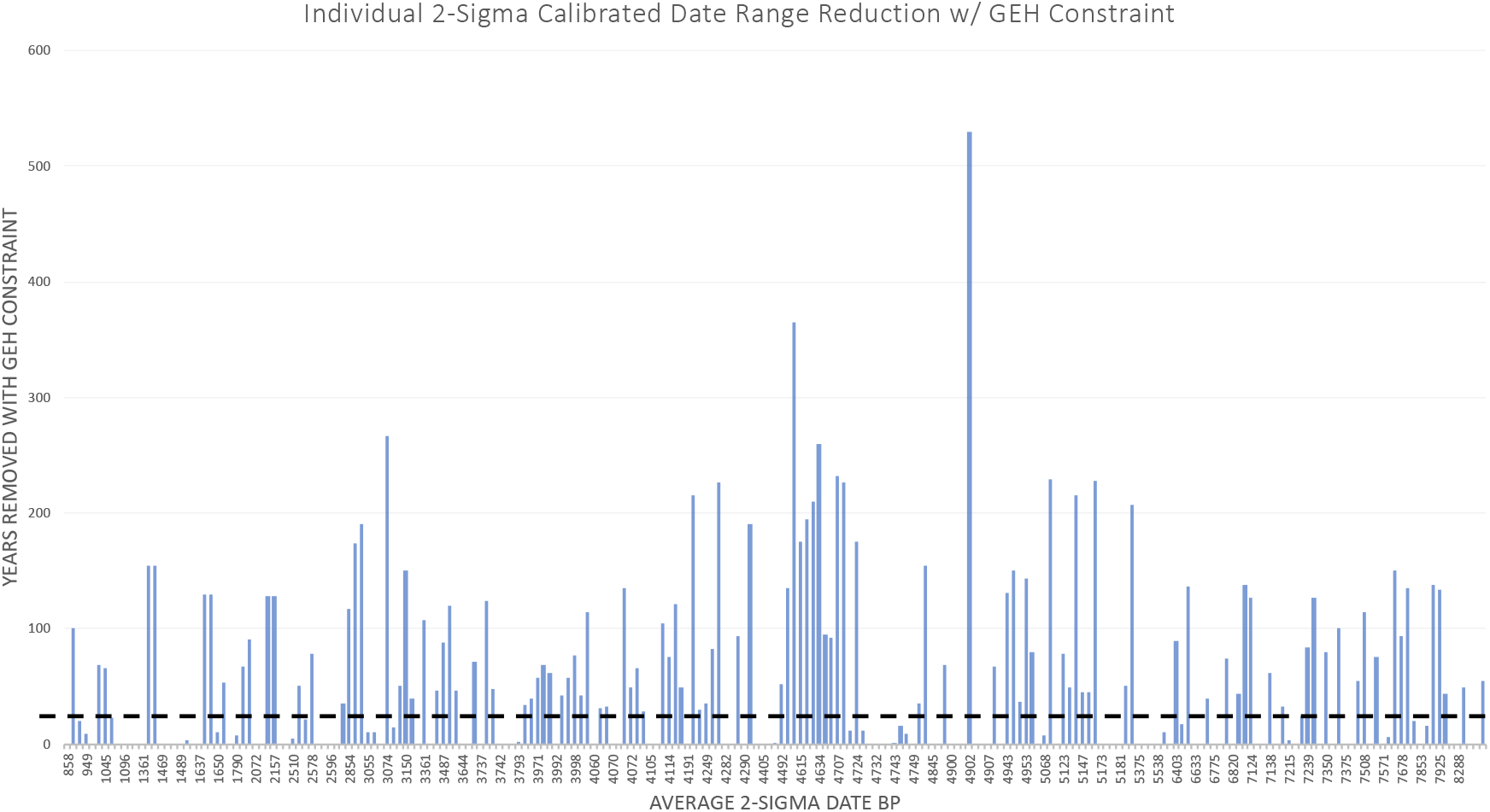
Difference between original 2 sigma calibrated date range and GH constrained range for all 219 individuals, ordered from most recent date BP to oldest.

One possible reason for large date range reductions is if different skeletal elements were radiocarbon dated for each individual in the relative pairs. Studies have demonstrated that different skeletal elements have different rates of remodeling and carbon uptake (Calcagnile et al., 2013; Cook et al., 2015; Hansen et al., 2017; Pinhasi et al., 2015); for example, a long bone (tibia, femur, etc.) remodels throughout an individual’s life and therefore regularly uptakes new carbon, whereas the otic capsule completes formation in utero and does not remodel during an individual’s lifetime. Thus, if a femur and otic capsule from the same individual are radiocarbon dated, two different dates may be generated, particularly in advanced-age individuals. This could potentially lead to discrepant date ranges for related individuals—if the otic capsule of an adult female who died giving childbirth was dated, while the femur of her daughter was used, there could in theory be a difference of more than 100 years. We therefore compiled data on which element of each individual was radiocarbon dated; information on which skeletal element was radiocarbon dated for each individual is provided in SM 4.1. Unfortunately, in many instances no information on which element was dated was available in the published literature. Additionally, if information was provided, it was sometimes imprecise or vague. An element might be listed as “petrous”, but with no information on if the otic capsule, cochlea, or ossicles were radiocarbon dated—these could generate earlier dates than the surrounding petrous pyramid or temporal bone.

We hypothesized that there would be a higher number of individuals above the 54.47 GH mean reduction for individuals in a related pair that had different skeletal elements C14 dated. Table 4 provides counts of whether the skeletal element dated for each individual was different or the same (or if no information was available) as their relative. While there does seem to be a higher proportion of individuals with reductions above the 54.47 year GH mean in instances where different elements were radiocarbon dated, this is not statistically significant (chi-square test; x^2^=1.96, p value= 0.3755, df=2; SM 4.4), suggesting that dating different elements of related pairs does not significantly impact the reductions made with GH constraints. This likely is due to the fact that despite different elements being dated, most relatives were relatively close in age (likely because few individuals in pre-modern societies reached advanced age), or that the different elements dated had similar bone remodeling/carbon uptake rates. Despite this, it is likely that some instances of large discrepancies can be explained by the C14 dating of different skeletal elements.

**Table 4.** Counts and percentage of whether the same or different skeletal elements were used to date an individual in a relative pair.

| **Comparison of skeletal element dated for each related pair** | **N individuals** | **% above 54.47 year mean GH reduction per element category** |
| --- | --- | --- |
| Different elements | 26 | 50.0 |
| Same element | 138 | 30.9 |
| No information available | 55 | 34.1 |

We next examined whether applying GH constraints could reveal larger patterns in the dataset. To explore if there were any periods which had a higher number of relatives that exceeded the mean reduction than others, we binned into 500-year intervals all 3965 published individuals in the database and the 219 dated individuals with genetically identified relatives. This allowed us to examine if periods with high numbers of individuals above the 54.47 year mean GH reduction were indicative of anomalies in the radiocarbon record (i.e. calibration curve issues, uncorrected marine reservoir effects, etc) at a particular date interval, or merely an artifact of sampling the database. Figures 9a and 9b qualitatively demonstrate that periods with a high number of individuals with a reduction above the 54.47 year mean roughly corresponds with the 500-year intervals that have been most densely sampled for aDNA. We performed a x^2^ test to test the null hypothesis that the number of individuals above the 54.47 year mean GH reductions per 500-year interval correlates with the total number of individuals sampled per 500-year interval. We found a x^2^ value of 57.43 (df=20), giving a p value= 1.76717E-05 (SM 4.5), rejecting the null hypothesis and suggesting that the number of individuals above the 54.47 mean GH reduction per 500-year interval is not simply due to the overall number of individuals sampled per 500-year interval. The most notable intervals were 7999-7500 BP, 7499-7000 BP, 4999-4500 BP, and 4499-4000 BP (Figure 9.C), which had residual values of 4.86, 5.07, 11.59, and 1.65, respectively (SM 4.5).

**Figure 9.**
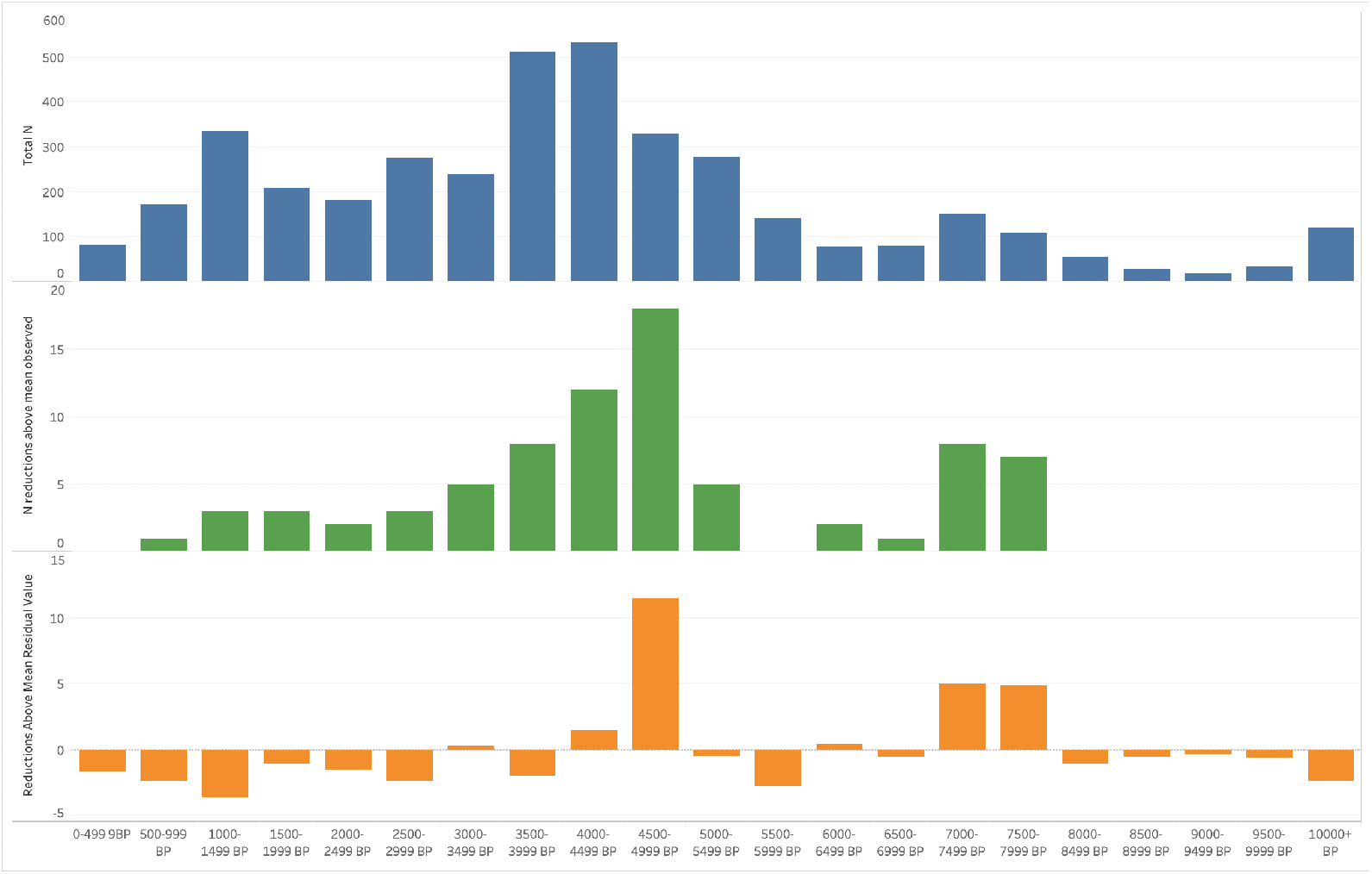
Data from the database and 219 individuals with relatives binned in 500-year intervals. A) #of individuals total in each 500-year interval. B) # of relatives per interval that exceed the 54.47-year mean GH reduction C) Residuals from x^2^ test.

The 500-year interval with the most individuals that had range reductions above the 54.47-year mean (n=18) was 4999-4500 calBP. The plotted radiocarbon distributions of these individuals demonstrate that plateaus along the radiocarbon curve during this period of time could account for the high number of reductions (Figs 10), which is also true for the other three 500-year intervals with the highest x^2^ residual values (Figs 11–13).

**Figure 10.**
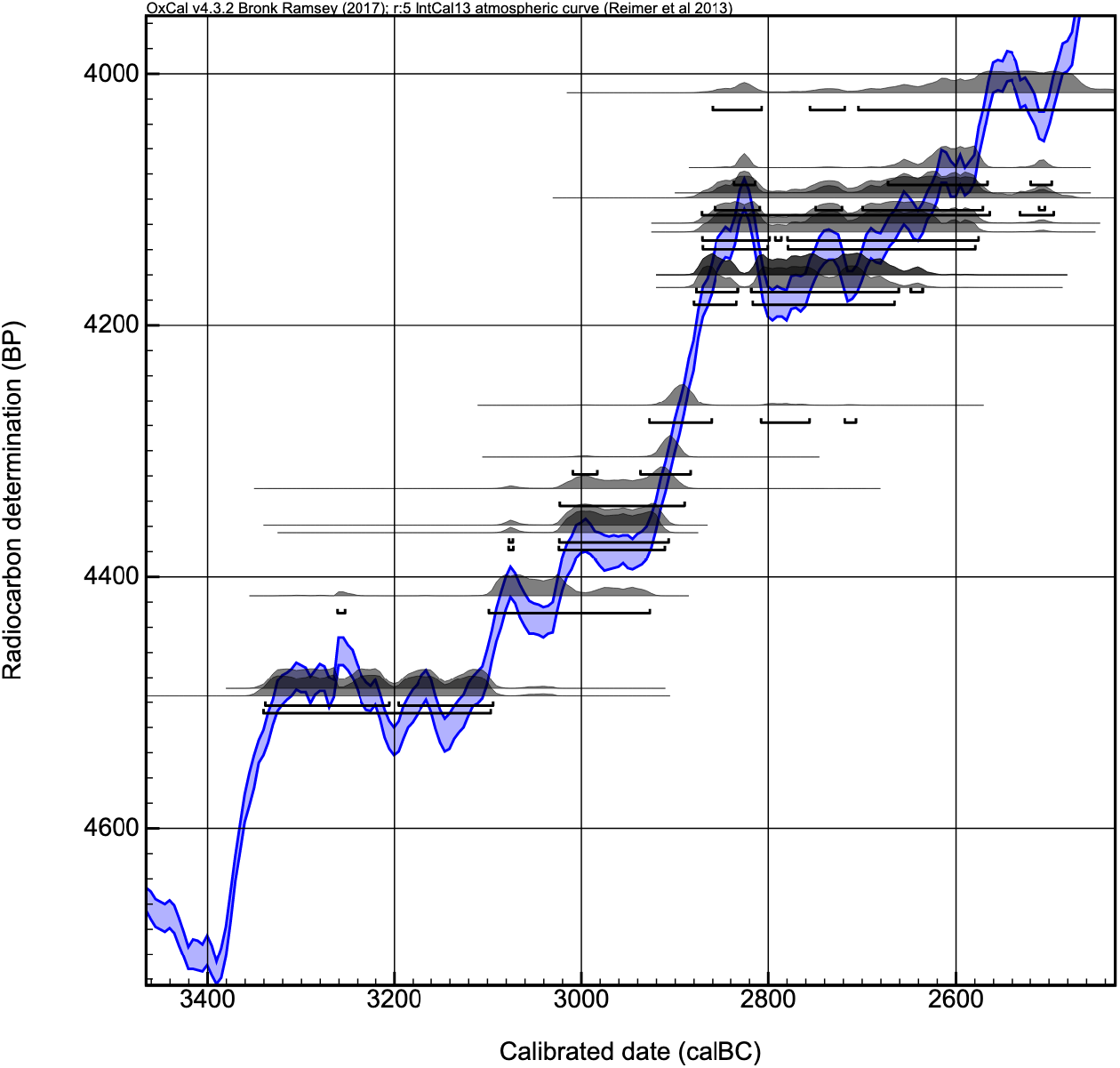
Original 2-sigma calBCE date ranges for individuals from the 4500-4999 calBP interval that had GH reductions about the 54.47 mean.

**Figure 11.**
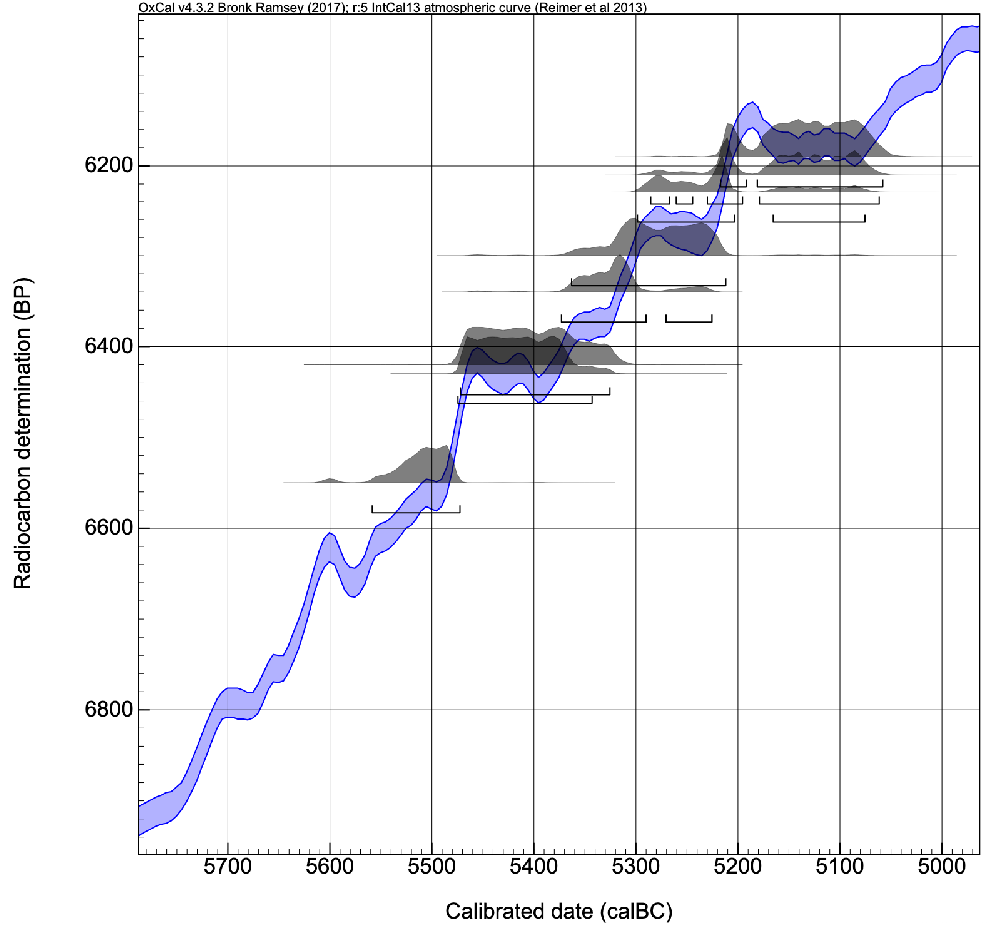
Original 2-sigma calBCE date ranges for individuals from the 7499-7000 calBP interval that had GH reductions about the 54.47 mean.

**Figure 12.**
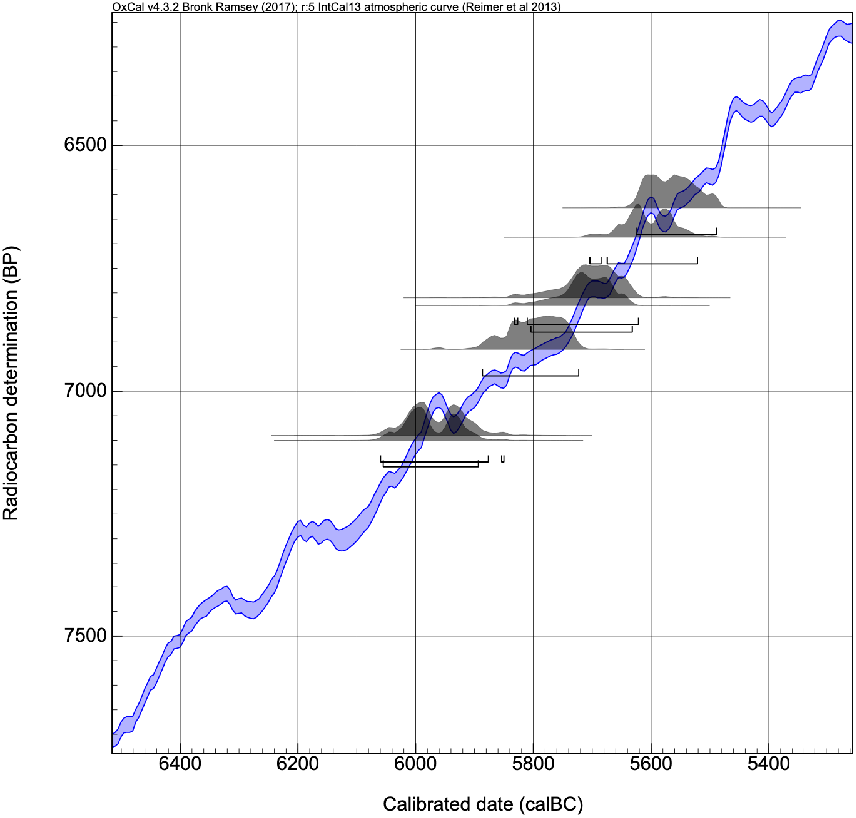
Original 2-sigma calBCE date ranges for individuals from the 7500-7999 calBP interval that had GH reductions about the 54.47 mean.

**Figure 13.**
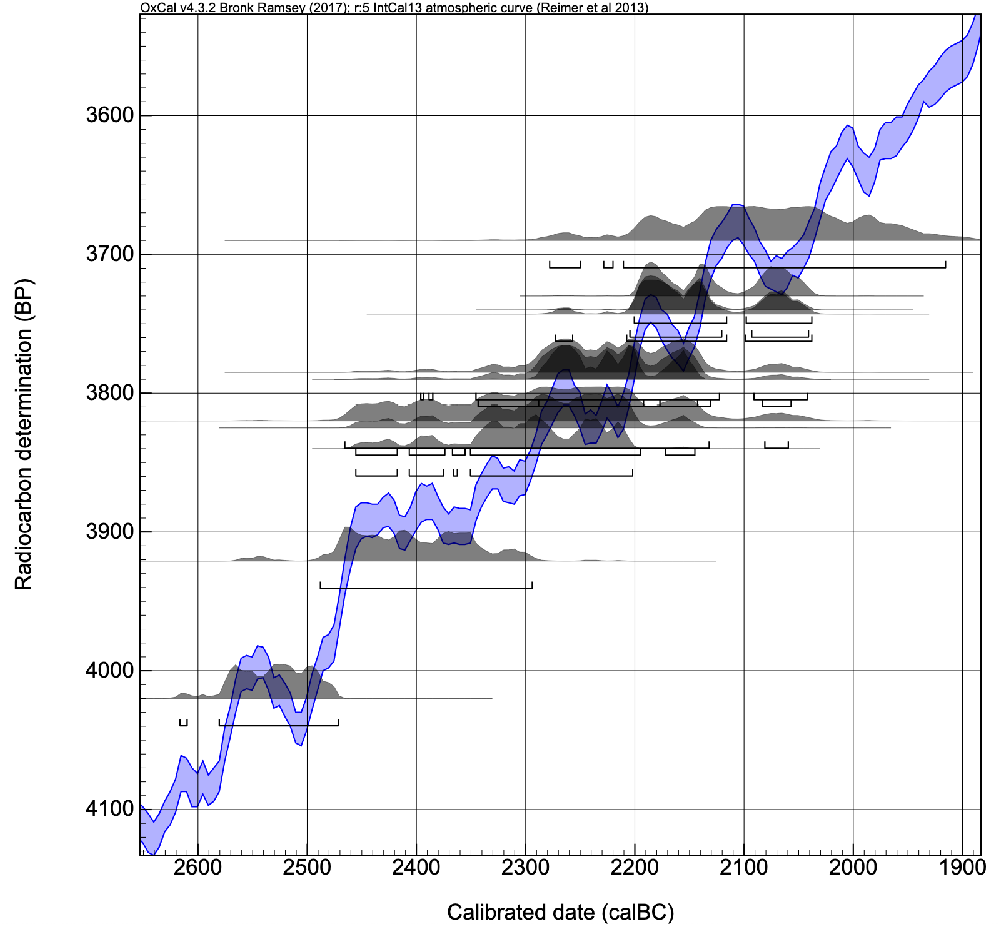
Original 2-sigma calBCE date ranges for individuals from the 4000-4499 BP calBP interval that had GH reductions about the 54.47 mean.

Individuals MX188 and MX190, 1^st^ degree relatives from Spreitenbach, Switzerland (Furtwangler et al. 2020), demonstrate how knowing genetic relatedness and applying GH constraints can reduce date ranges and help correct for radiocarbon plateaus. MX 190’s 2-sigma range falls on a curve plateau and is much larger than MX 188’s (Figure 14; 2861-2342 calBCE, ETH-19935 and 2495-2399 calBCE, BE-7995.1.1, respectively). However, because these individuals are known to be 1^st^ degree relatives the 29 year GH constraint could be applied, reducing MX 190’s range by 365 years (to 2524-2370 calBCE, SM 4.1).

**Figure 14.**
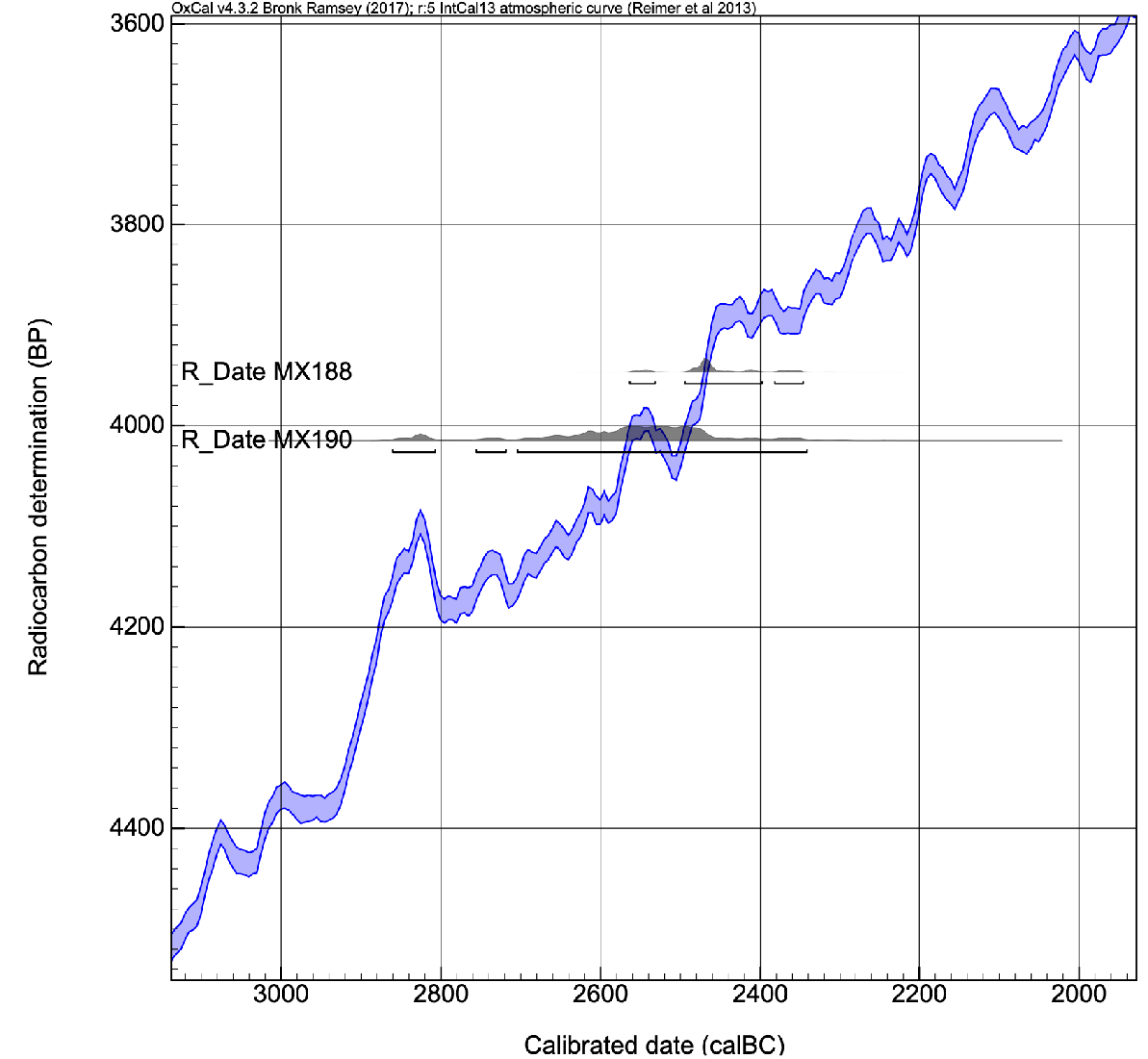
Original 2-sigma calBCE date ranges for MX 188 and MX 190.

### 3.6 Building Bayesian Models

The range tightening described above is a “manual” method for constraining the tail ends of radiocarbon date distributions using estimates for the number of years that can separate the dates of death of genetic relatives. While date ranges can be constrained with this manual method, we also tested how other statistical modeling could refine the date ranges. Bayesian analysis to increase precision in a series of radiocarbon dates has become standard practice amongst archaeologists (Bronk Ramsey, 2009; Taylor and Bar-Yosef, 2014). Thus, we examined how effective knowledge of genetic relatedness and date of death estimates are as priors to refine radiocarbon dates.

To test this, we started by importing the raw calibrated date probability distributions for I2600 and I2457 from OxCal 4.3 (data provided in SM 5). We next sorted the GH DOD values into 5-year intervals and produced a probability distribution. The raw data were smoothed to give estimates of DOD by year (SM 5). The posterior joint distribution of the datasets was then computed. Figures 15 and 16 are the marginal estimates of these distributions for the father and daughter. Due to a plateau in the radiocarbon curve, the date distributions for the father and daughter are bimodal with an additional, lower probability “peak” (demonstrated with the purple curve). For each, building in the relative information significantly reduces the probability of one of the original probability peaks. And while the distribution for I2600 essentially remains bimodal, the most likely probability for both I2600 and I2457 is between 2100-2000 BCE. This result demonstrates that building the constraints in to statistical modeling can help refine date ranges. Future work building these constraints into Bayesian modeling available in OxCal could provide additional refinements.

**Figure 15.**
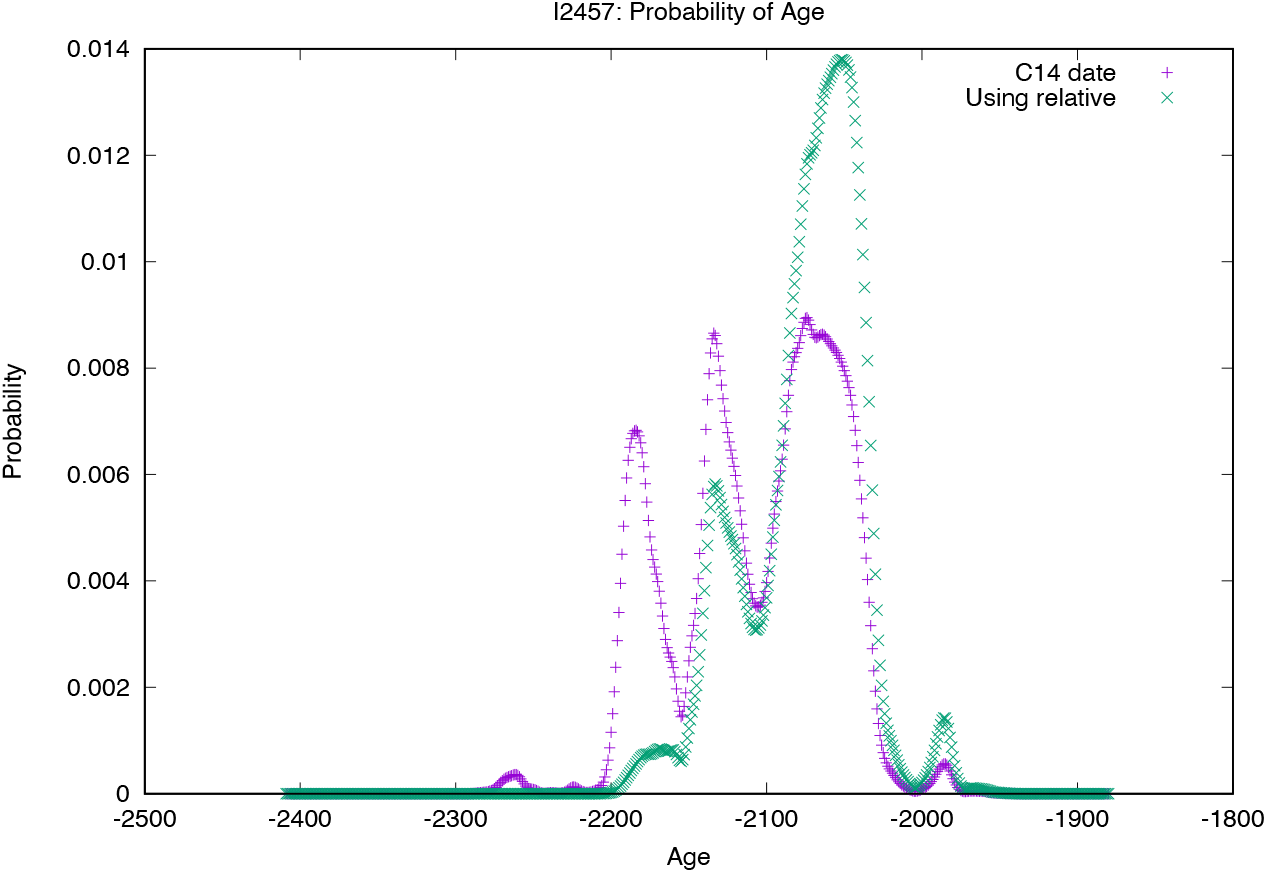
Joint probability distribution for I2457. Original AMS date probability distribution in purple, new joint distribution in green.

**Figure 16.**
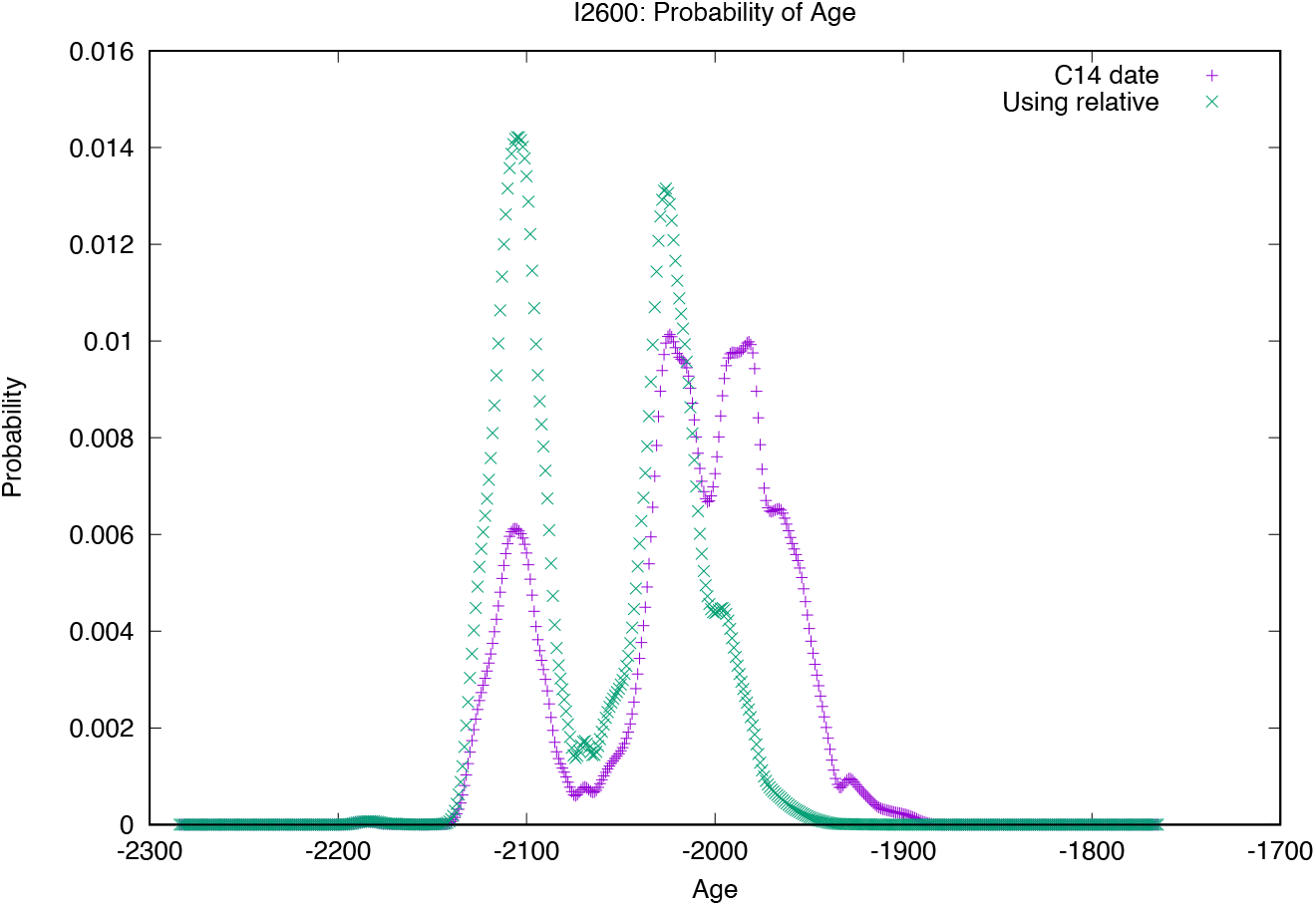
Joint probability distribution for I2600. Original AMS date probability distribution in purple, new joint distribution in green.

### 3.7. Summary

In sum, knowledge of genetic relatedness can be used to constrain radiocarbon date distributions, either by applying DOD separations to the tail ends of the distributions, or through Bayesian modeling. These refinements are not universally applicable; related pairs often have date distributions that overlap, sometimes almost entirely, limiting the extent to which DOD estimates can refine date ranges. Yet, overlap is what *should* be expected; related individuals should not have large date separations. Date distributions of related individuals that do not overlap could reveal an error in radiocarbon dating (such as I2600 and I2457) or genetic analysis, or other issues, such as an uncorrected marine reservoir effect. In other words, the more substantially DOD separation estimates can constrain C14 date ranges, the more likely a significant issue exists in dating for any of a variety of reasons (unaccounted marine reservoir effect, curve plateau, etc).

## 4. Discussion

Combining previously independent lines of data—knowledge of genetic relatedness derived from ancient DNA; biological and estimated DOD separations for relatives; and radiocarbon dates—creates potential benefits for researchers examining the ancient past. Perhaps the most apparent is evaluation of data generated through disparate methods. As discussed above, relatedness often confirms radiocarbon dates (and vice-versa, Saag et al., 2019:5). Using genetic relatedness and DOD separation estimates to evaluate radiocarbon dates can also help attend to some of the most common pitfalls in radiocarbon dating. According to Taylor and Bar-Yosef (2014:132), “the most common reason why C14 dating evidence is considered to be anomalous can be traced to failures to clearly establish and document the physical relationship between a C14 dated sample and a specific targeted event or cultural expression.” Somewhat counterintuitively, incorporating genetic relationship DOD-separations addresses Taylor and Bar-Yosef’s concerns by circumventing taphonomic processes. Instead of focusing on potential confounding factors of when individuals were buried, removed, reburied, etc., date ranges are examined with an independent line of evidence that is not prone to contamination issues associated with taphonomic processes and archaeological context.

Analyzing the radiocarbon record with knowledge of genetic relatives also provides archaeologists an opportunity to move beyond traditional interpretations of radiocarbon dates. C14 dating of skeletal remains typically provides an estimate of when an individual died (although it could also reflect when a particular element ceased carbon uptake during an individual’s lifetime, as discussed above). The combined and constrained date ranges discussed above provide minimum and maximum boundaries of when two related individuals died; therefore, the overlap of the two ranges likely contains the plausible period of time when the individuals were *alive* together. It is entirely possible, of course, that one individual died at the minimum boundary and the other at the maximum. In such instances the two related individuals would have no lifetime overlap. Even in such cases, considering the combined C14 ranges of related individuals can turn archaeological thinking away from incipient or terminal dates of archaeological periods, but instead toward changes that happened during lifetimes.

Considering lifespan ranges also helps elucidate cultural plasticity, and reveal the arbitrariness of archaeological boundaries. Archaeologists had initially suspected the English Bell Beaker father-daughter pair were part of separate archaeological cultures due to their initial dates. Knowing that these two individuals were related not only helped in identifying an error in the initial radiocarbon dates, but also speaks to the subjective nature of chronological and cultural boundaries archaeologists establish, which were of no consequence for the father-daughter pair.

The approaches outlined above represent only a small number of applications for how knowledge of genetic relatedness can help with radiocarbon dating. The potential for further applicability needs to be explored; one promising application could be the use of extended families for radiocarbon curve “wiggle matching.”

## 5. Conclusion

This research is a first step in combining two discrete analytical methods to add refinement to interpretation of the archaeological record and is meant to demonstrate that knowledge of genetic relatedness can be used to augment radiocarbon dating. As ancient DNA databases continue to grow, and more relatives are identified and radiocarbon dated, researchers will likely feel compelled to refine GH and DOD estimates as they see fit, as some have already done (Kennett et al., 2017; Saag et al., 2019). Levels of social organization (e.g. hunter-gatherer vs. agriculturalist), age of skeletons (adult vs. juvenile), and lifespan estimate could also all be incorporated into estimates. Additionally, once enough related individuals are identified and dated, specific regions, sub-regions, or even sites can be examined for anomalies in the associated radiocarbon records.

Ancient DNA innovations are providing archaeologists with unprecedented insight into the past. As ancient DNA becomes increasingly integral to archaeological studies, researchers should explore novel applications of genetic data to archaeological studies. This paper used ancient DNA to identify radiocarbon outliers, refine date distribution ranges for related pairs, and delineate potential issues unaccounted for in the radiocarbon record of particular eras and locales. Such studies should help integrate the two fields and move ancient DNA and archaeology forward together into the next era of research on the human past.

## Supporting information

Supplemental Data References

## Declaration of Competing Interests

None

## Acknowledgements

This work was funded by NIH grant GM100233, the Paul Allen Foundation, John Templeton Foundation (grant number 6122), and David Reich is an Investigator of the Howard Hughes Medical Institute. We thank Melissa Gymrek for providing assistance with familinx data from the Kaplanis et al. 2018 study, Greg Hodgins for reviewing an early draft of the paper, Iain Mathieson and Vagheesh Narasimhan for statistical assistance, and members of our laboratory for feedback on the study during its development.

## Notes

https://familinx.org

http://www.rootsweb.ancestry.com/~okcaddo/cemeteries/alfalfa.txt

http://www.rootsweb.ancestry.com/~okcaddo/cemeteries/bethelmn.txt

http://www.genealogy.com/ftm/n/e/y/Brenda-K-Neyra/GENE11-0001.html

https://www.findagrave.com/cgi-bin/fg.cgi?page=vcsr&GSsr=1&GSvcid=292671&

https://wc.rootsweb.ancestry.com/cgi-bin/igm.cgi?op=SRCH&db=klea&surname=A

https://reich.hms.harvard.edu/downloadable-genotypes-present-day-and-ancient-dna-data-compiled-published-papers

### Competing Interest Statement

The authors have declared no competing interest.

