## Supplemental Figures for "Combining Ancient DNA and Radiocarbon Dating Data to Increase Chronological Precision"

#### Biological Maximum DOD Separation: Parents-Offspring

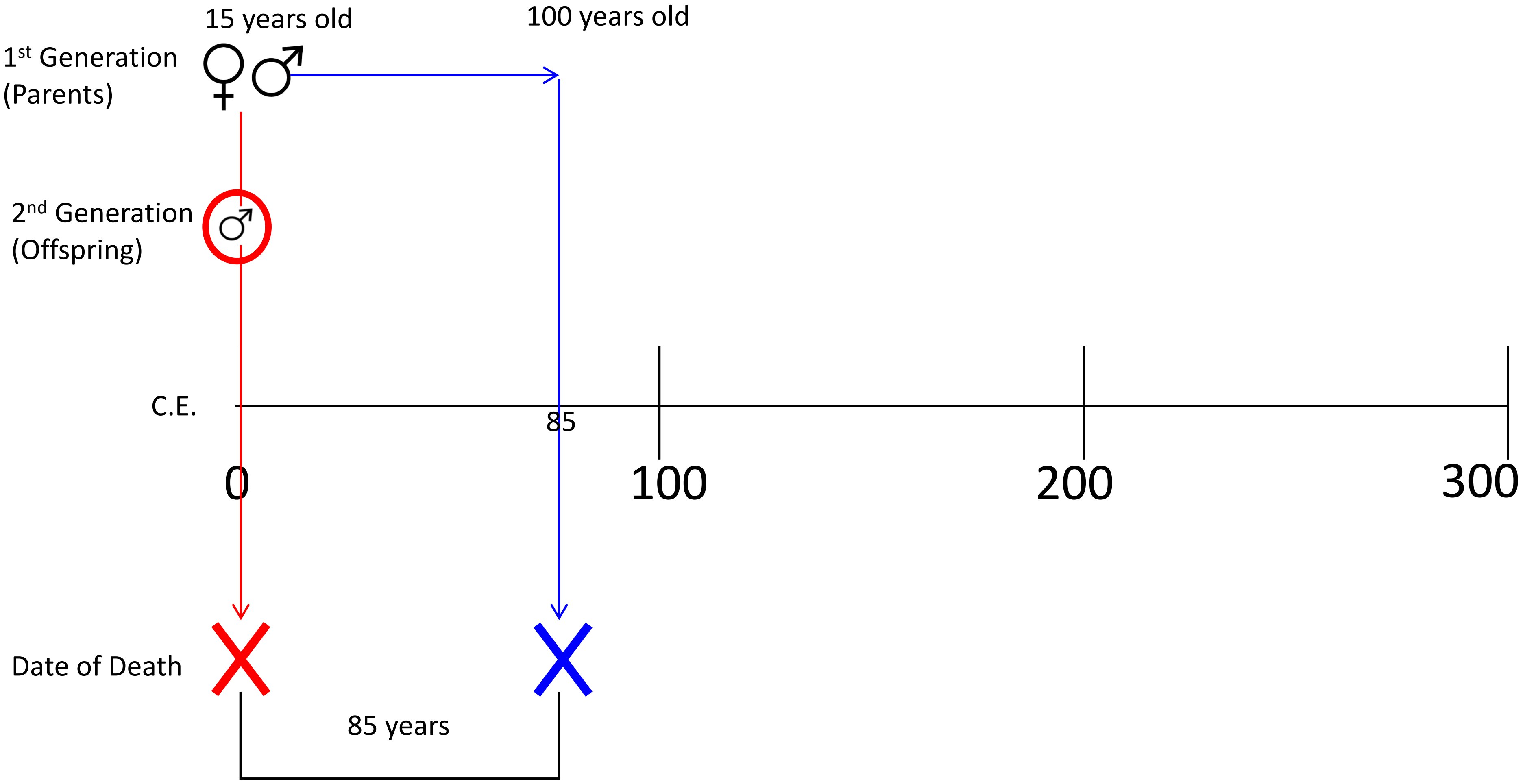

#### Biological Maximum DOD Separation: Parents-Offspring Alternative

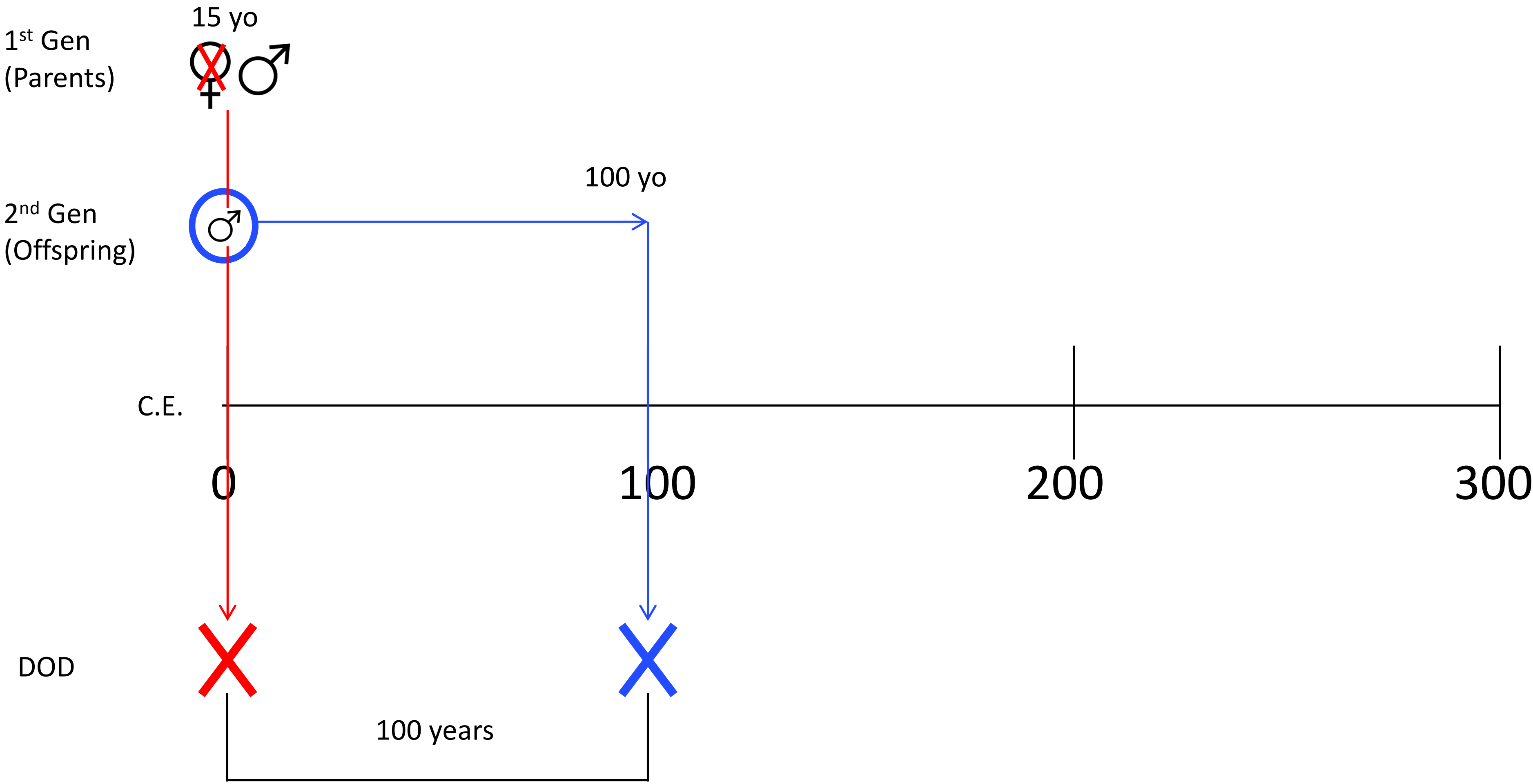

### Biological Maximum DOD Separation: Siblings

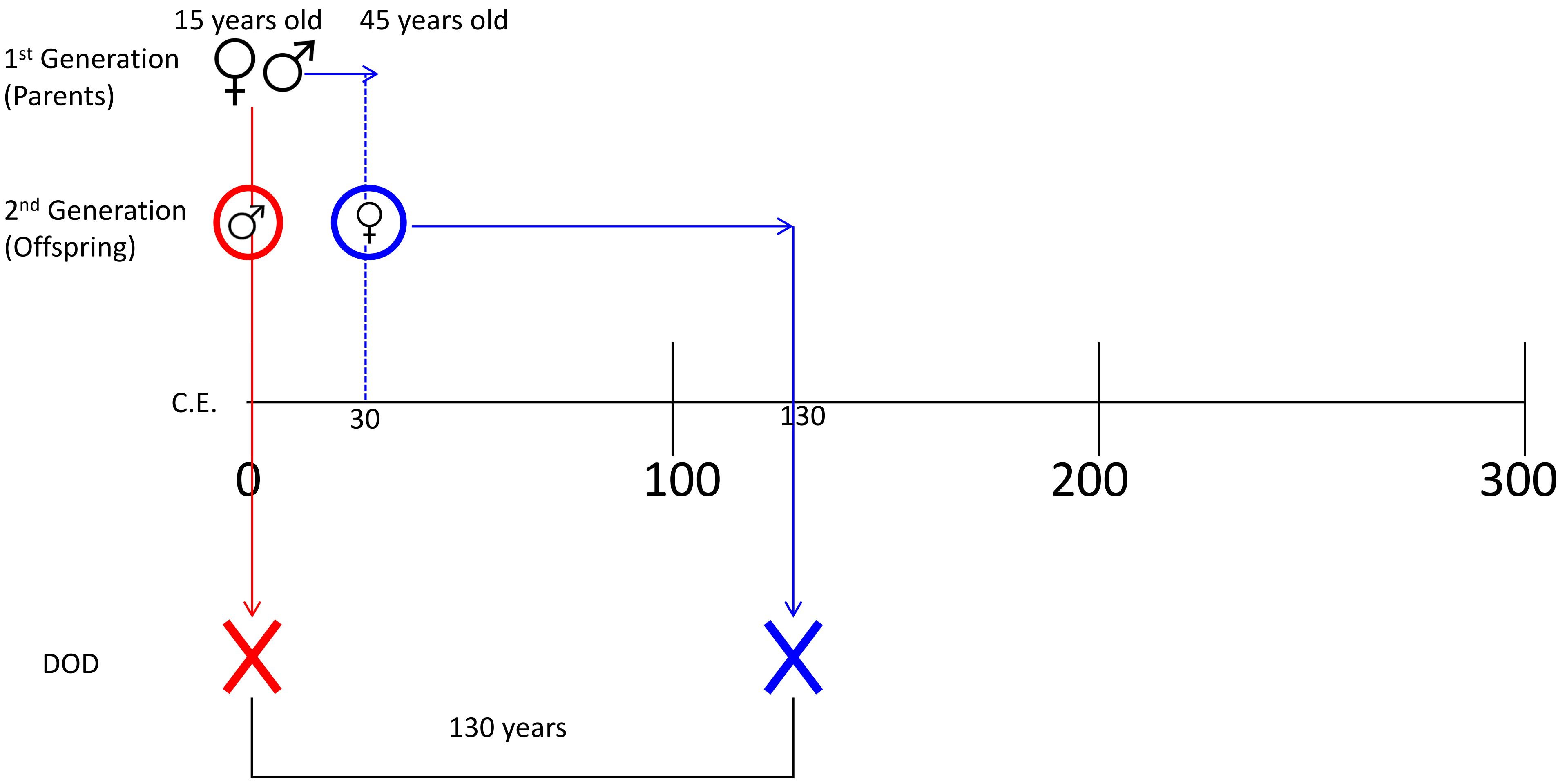

### Biological Maximum DOD Separation: Grandparent-Grandchild

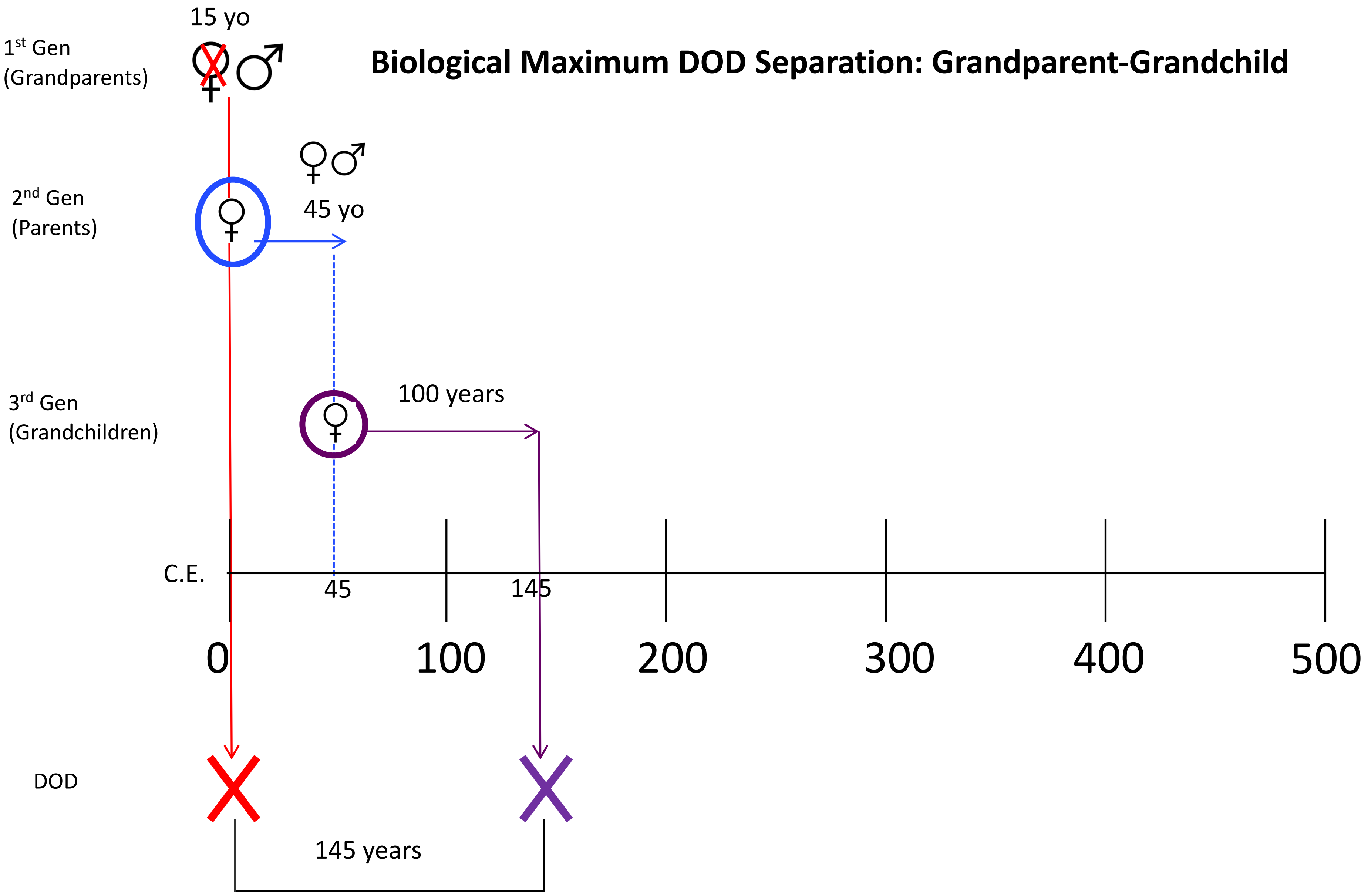

### Biological Maximum DOD Separation: Grandparent-Grandchild Alternative

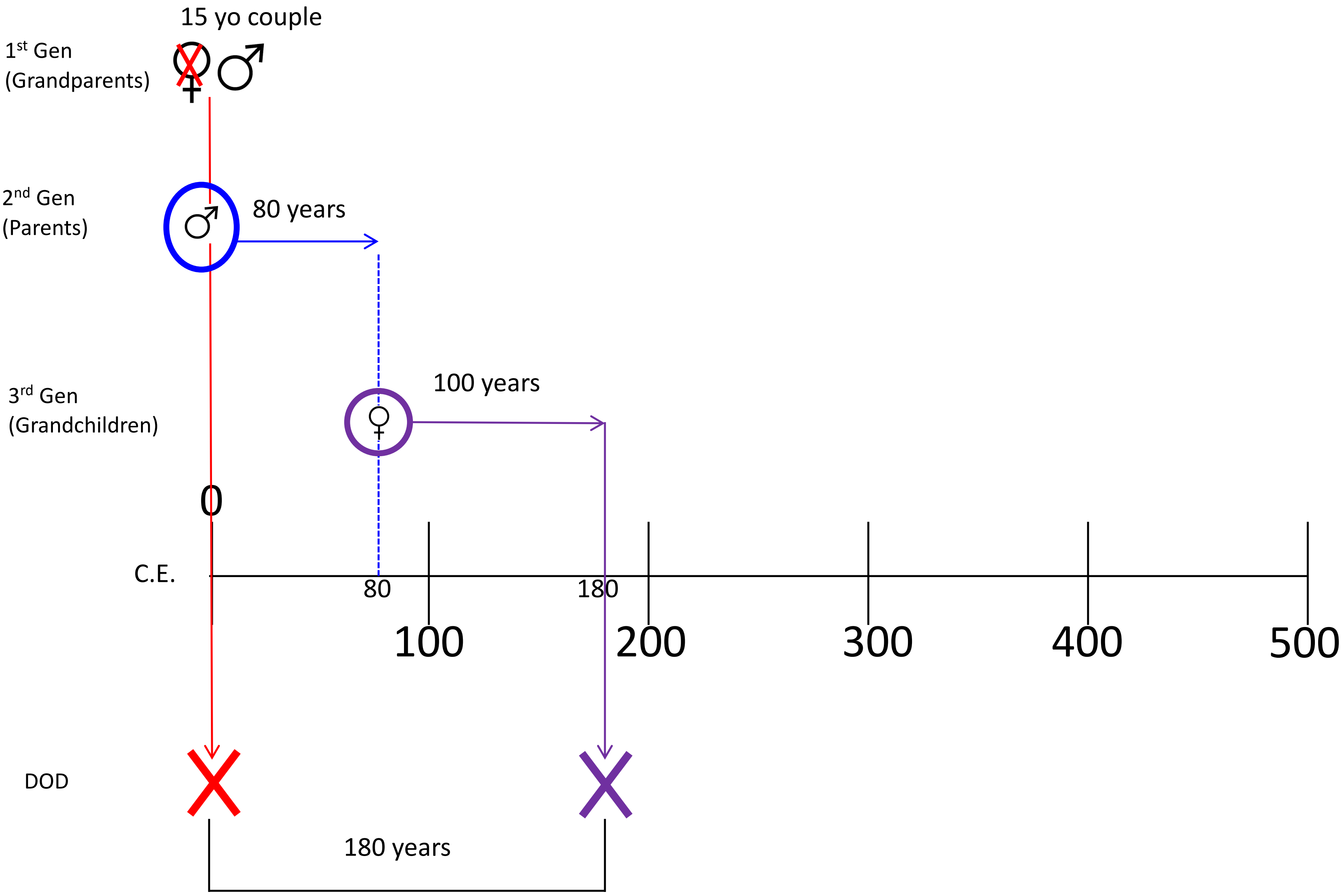

### Biological Maximum DOD Separation: Cousins

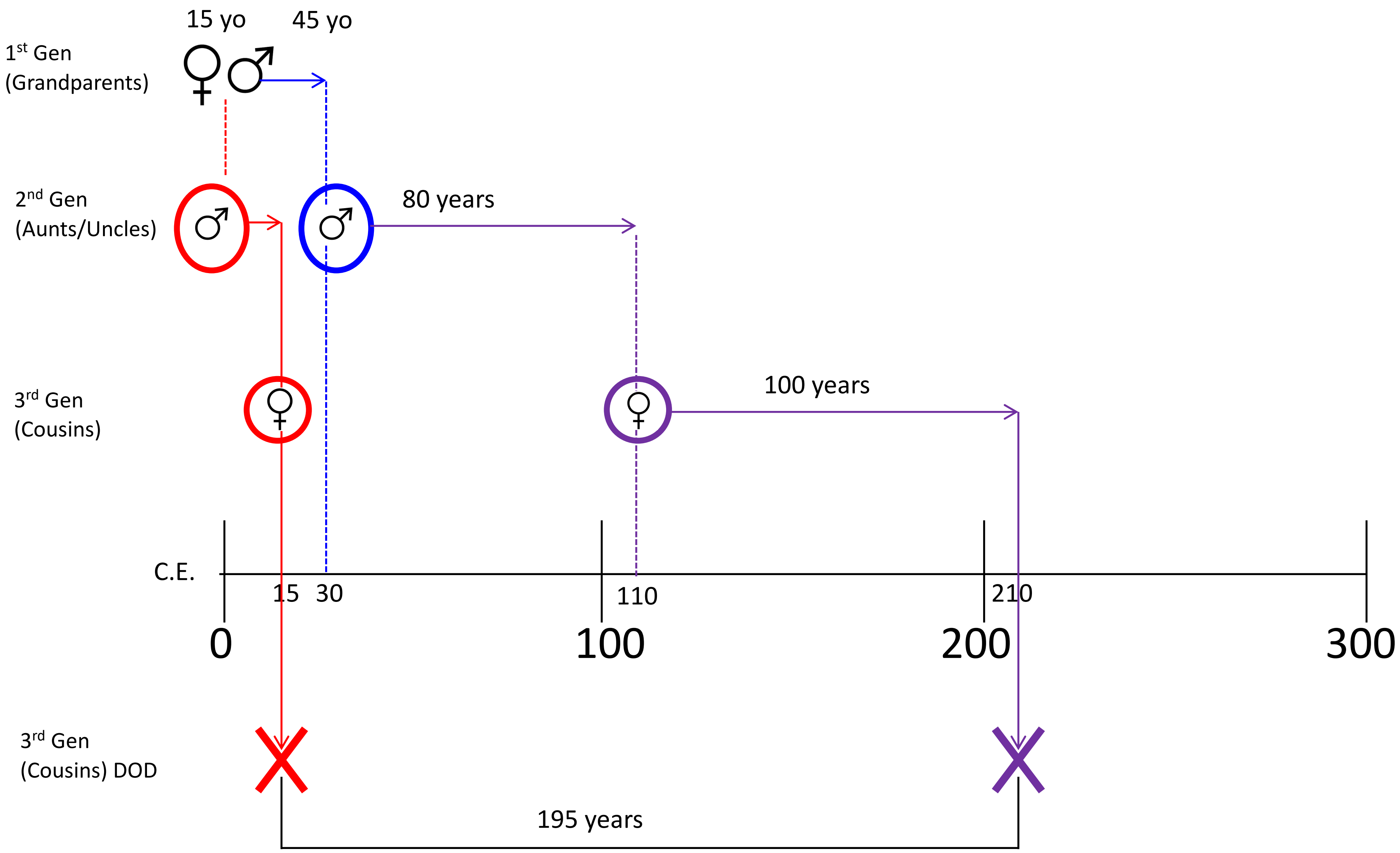

### Biological Maximum DOD Separation: Aunts/Uncles

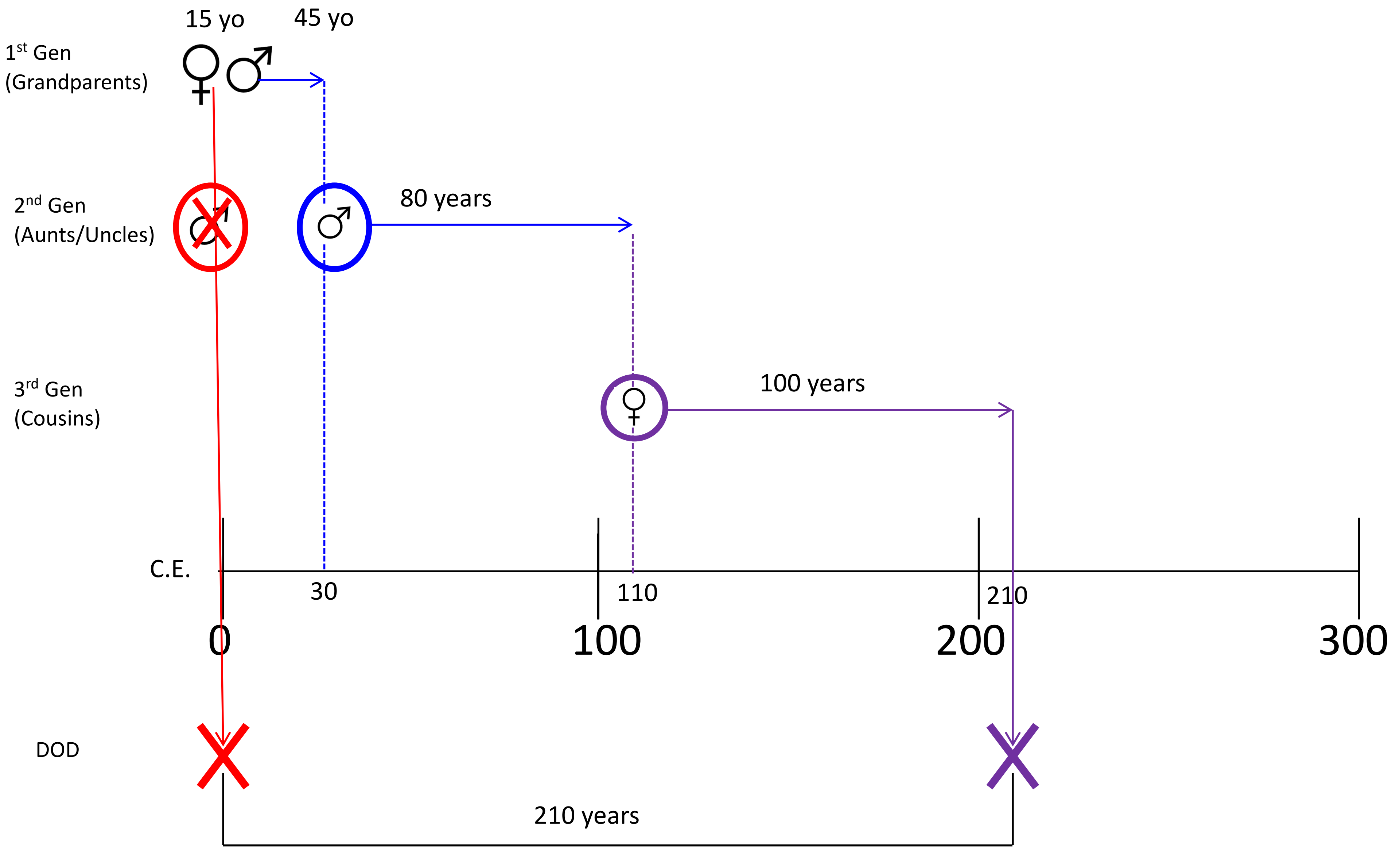
