## Supplemental Data References for "Combining Ancient DNA and Radiocarbon Dating Data to Increase Chronological Precision"

de Barros Damgaard, P., Martiniano, R., Kamm, J., Moreno-Mayar, J.V., Kroonen, G., Peyrot, M., Barjamovic, G., Rasmussen, S., Zacho, C., Baimukhanov, N., Zaibert, V., Merz, V., Biddanda, A., Merz, I., Loman, V., Evdokimov, V., Usmanova, E., Hemphill, B., Seguin-Orlando, A., Yediay, F.E., Ullah, I., Sjögren, K.-G., Iversen, K.H., Choin, J., de la Fuente, C., Ilardo, M., Schroeder, H., Moiseyev, V., Gromov, A., Polyakov, A., Omura, S., Senyurt, S.Y., Ahmad, H., McKenzie, C., Margaryan, A., Hameed, A., Samad, A., Gul, N., Khokhar, M.H., Goriunova, O.I., Bazaliiskii, V.I., Novembre, J., Weber, A.W., Orlando, L., Allentoft, M.E., Nielsen, R., Kristiansen, K., Sikora, M., Outram, A.K., Durbin, R., Willerslev, E., 2018. The first horse herders and the impact of early Bronze Age steppe expansions into Asia. Science 360, eaar7711. https://doi.org/10.1126/science.aar7711

Fernandes, D.M., Mittnik, A., Olalde, I., Lazaridis, I., Cheronet, O., Rohland, N., Mallick, S., Bernardos, R., Broomandkhoshbacht, N., Carlsson, J., Culleton, B.J., Ferry, M., Gamarra, B., Lari, M., Mah, M., Michel, M., Modi, A., Novak, M., Oppenheimer, J., Sirak, K.A., Stewardson, K., Mandl, K., Schattke, C., Özdoğan, K.T., Lucci, M., Gasperetti, G., Candilio, F., Salis, G., Vai, S., Camarós, E., Calò, C., Catalano, G., Cueto, M., Forgia, V., Lozano, M., Marini, E., Micheletti, M., Miccichè, R.M., Palombo, M.R., Ramis, D., Schimmenti, V., Sureda, P., Teira, L., Teschler-Nicola, M., Kennett, D.J., Lalueza-Fox, C., Patterson, N., Sineo, L., Coppa, A., Caramelli, D., Pinhasi, R., Reich, D., 2020. The spread of steppe and Iranian-related ancestry in the islands of the western Mediterranean. Nature Ecology & Evolution 4, 334–345. https://doi.org/10.1038/s41559-020-1102-0

Flegontov, P., Altınışık, N.E., Changmai, P., Rohland, N., Mallick, S., Adamski, N., Bolnick, D.A., Broomandkhoshbacht, N., Candilio, F., Culleton, B.J., Flegontova, O., Friesen, T.M., Jeong, C., Harper, T.K., Keating, D., Kennett, D.J., Kim, A.M., Lamnidis, T.C., Lawson, A.M., Olalde, I., Oppenheimer, J., Potter, B.A., Raff, J., Sattler, R.A., Skoglund, P., Stewardson, K., Vajda, E.J., Vasilyev, S., Veselovskaya, E., Hayes, M.G., O’Rourke, D.H., Krause, J., Pinhasi, R., Reich, D., Schiffels, S., 2019. Palaeo-Eskimo genetic ancestry and the peopling of Chukotka and North America. Nature 570, 236–240. https://doi.org/10.1038/s41586-019-1251-y

Furtwängler, A., Rohrlach, A.B., Lamnidis, T.C., Papac, L., Neumann, G.U., Siebke, I., Reiter, E., Steuri, N., Hald, J., Denaire, A., Schnitzler, B., Wahl, J., Ramstein, M., Schuenemann, V.J., Stockhammer, P.W., Hafner, A., Lösch, S., Haak, W., Schiffels, S., Krause, J., 2020. Ancient genomes reveal social and genetic structure of Late Neolithic Switzerland. Nature Communications 11, 1915. https://doi.org/10.1038/s41467-020-15560-x

Haber, M., Doumet-Serhal, C., Scheib, C., Xue, Y., Danecek, P., Mezzavilla, M., Youhanna, S., Martiniano, R., Prado-Martinez, J., Szpak, M., Matisoo-Smith, E., Schutkowski, H., Mikulski, R., Zalloua, P., Kivisild, T., Tyler-Smith, C., 2017. Continuity and Admixture in the Last Five Millennia of Levantine History from Ancient Canaanite and Present-Day Lebanese Genome Sequences. The American Journal of Human Genetics 101, 274–282. https://doi.org/10.1016/j.ajhg.2017.06.013

Järve, M., Saag, Lehti, Scheib, C.L., Pathak, A.K., Montinaro, F., Pagani, L., Flores, R., Guellil, M., Saag, Lauri, Tambets, K., Kushniarevich, A., Solnik, A., Varul, L., Zadnikov, S., Petrauskas, O., Avramenko, M., Magomedov, B., Didenko, S., Toshev, G., Bruyako, I., Grechko, D., Okatenko, V., Gorbenko, K., Smyrnov, O., Heiko, A., Reida, R., Sapiehin, S., Sirotin, S., Tairov, A., Beisenov, A., Starodubtsev, M., Vasilev, V., Nechvaloda, A., Atabiev, B., Litvinov, S., Ekomasova, N., Dzhaubermezov, M., Voroniatov, S., Utevska, O., Shramko, I., Khusnutdinova, E., Metspalu, M., Savelev, N., Kriiska, A., Kivisild, T., Villems, R., 2019. Shifts in the Genetic Landscape of the Western Eurasian Steppe Associated with the Beginning and End of the Scythian Dominance. Current Biology 29, 2430-2441.e10. https://doi.org/10.1016/j.cub.2019.06.019

Kennett, D.J., Plog, S., George, R.J., Culleton, B.J., Watson, A.S., Skoglund, P., Rohland, N., Mallick, S., Stewardson, K., Kistler, L., LeBlanc, S.A., Whiteley, P.M., Reich, D., Perry, G.H., 2017. Archaeogenomic evidence reveals prehistoric matrilineal dynasty. Nature Communications 8, 14115. https://doi.org/10.1038/ncomms14115

Krzewińska, M., Kjellström, A., Günther, T., Hedenstierna-Jonson, C., Zachrisson, T., Omrak, A., Yaka, R., Kılınç, G.M., Somel, M., Sobrado, V., Evans, J., Knipper, C., Jakobsson, M., Storå, J., Götherström, A., 2018. Genomic and Strontium Isotope Variation Reveal Immigration Patterns in a Viking Age Town. Current Biology 28, 2730-2738.e10. https://doi.org/10.1016/j.cub.2018.06.053

Linderholm, A., Kılınç, G.M., Szczepanek, A., Włodarczak, P., Jarosz, P., Belka, Z., Dopieralska, J., Werens, K., Górski, J., Mazurek, M., Hozer, M., Rybicka, M., Ostrowski, M., Bagińska, J., Koman, W., Rodríguez-Varela, R., Storå, J., Götherström, A., Krzewińska, M., 2020. Corded Ware cultural complexity uncovered using genomic and isotopic analysis from south-eastern Poland. Scientific Reports 10, 6885. https://doi.org/10.1038/s41598-020-63138-w

Lipson, M., Ribot, I., Mallick, S., Rohland, N., Olalde, I., Adamski, N., Broomandkhoshbacht, N., Lawson, A.M., López, S., Oppenheimer, J., Stewardson, K., Asombang, R.N., Bocherens, H., Bradman, N., Culleton, B.J., Cornelissen, E., Crevecoeur, I., de Maret, P., Fomine, F.L.M., Lavachery, P., Mindzie, C.M., Orban, R., Sawchuk, E., Semal, P., Thomas, M.G., Van Neer, W., Veeramah, K.R., Kennett, D.J., Patterson, N., Hellenthal, G., Lalueza-Fox, C., MacEachern, S., Prendergast, M.E., Reich, D., 2020. Ancient West African foragers in the context of African population history. Nature 577, 665–670. https://doi.org/10.1038/s41586-020-1929-1

Marcus, J.H., Posth, C., Ringbauer, H., Lai, L., Skeates, R., Sidore, C., Beckett, J., Furtwängler, A., Olivieri, A., Chiang, C.W.K., Al-Asadi, H., Dey, K., Joseph, T.A., Liu, C.-C., Der Sarkissian, C., Radzevičiūtė, R., Michel, M., Gradoli, M.G., Marongiu, P., Rubino, S., Mazzarello, V., Rovina, D., La Fragola, A., Serra, R.M., Bandiera, P., Bianucci, R., Pompianu, E., Murgia, C., Guirguis, M., Orquin, R.P., Tuross, N., van Dommelen, P., Haak, W., Reich, D., Schlessinger, D., Cucca, F., Krause, J., Novembre, J., 2020. Genetic history from the Middle Neolithic to present on the Mediterranean island of Sardinia. Nature Communications 11, 939. https://doi.org/10.1038/s41467-020-14523-6

Mathieson, I., Alpaslan-Roodenberg, S., Posth, C., Szécsényi-Nagy, A., Rohland, N., Mallick, S., Olalde, I., Broomandkhoshbacht, N., Candilio, F., Cheronet, O., Fernandes, D., Ferry, M., Gamarra, B., Fortes, G.G., Haak, W., Harney, E., Jones, E., Keating, D., Krause-Kyora, B., Kucukkalipci, I., Michel, M., Mittnik, A., Nägele, K., Novak, M., Oppenheimer, J., Patterson, N., Pfrengle, S., Sirak, K., Stewardson, K., Vai, S., Alexandrov, S., Alt, K.W., Andreescu, R., Antonović, D., Ash, A., Atanassova, N., Bacvarov, K., Gusztáv, M.B., Bocherens, H., Bolus, M., Boroneanţ, A., Boyadzhiev, Y., Budnik, A., Burmaz, J., Chohadzhiev, S., Conard, N.J., Cottiaux, R., Čuka, M., Cupillard, C., Drucker, D.G., Elenski, N., Francken, M., Galabova, B., Ganetsovski, G., Gély, B., Hajdu, T., Handzhyiska, V., Harvati, K., Higham, T., Iliev, S., Janković, I., Karavanić, I., Kennett, D.J., Komšo, D., Kozak, A., Labuda, D., Lari, M., Lazar, C., Leppek, M., Leshtakov, K., Vetro, D.L., Los, D., Lozanov, I., Malina, M., Martini, F., McSweeney, K., Meller, H., Menđušić, M., Mirea, P., Moiseyev, V., Petrova, V., Price, T.D., Simalcsik, A., Sineo, L., Šlaus, M., Slavchev, V., Stanev, P., Starović, A., Szeniczey, T., Talamo, S., Teschler-Nicola, M., Thevenet, C., Valchev, I., Valentin, F., Vasilyev, S., Veljanovska, F., Venelinova, S., Veselovskaya, E., Viola, B., Virag, C., Zaninović, J., Zäuner, S., Stockhammer, P.W., Catalano, G., Krauß, R., Caramelli, D., Zariņa, G., Gaydarska, B., Lillie, M., Nikitin, A.G., Potekhina, I., Papathanasiou, A., Borić, D., Bonsall, C., Krause, J., Pinhasi, R., Reich, D., 2018. The genomic history of southeastern Europe. Nature 555, 197–203. https://doi.org/10.1038/nature25778

Mathieson, I., Lazaridis, I., Rohland, N., Mallick, S., Patterson, N., Roodenberg, S.A., Harney, E., Stewardson, K., Fernandes, D., Novak, M., Sirak, K., Gamba, C., Jones, E.R., Llamas, B., Dryomov, S., Pickrell, J., Arsuaga, J.L., de Castro, J.M.B., Carbonell, E., Gerritsen, F., Khokhlov, A., Kuznetsov, P., Lozano, M., Meller, H., Mochalov, O., Moiseyev, V., Guerra, M.A.R., Roodenberg, J., Vergès, J.M., Krause, J., Cooper, A., Alt, K.W., Brown, D., Anthony, D., Lalueza-Fox, C., Haak, W., Pinhasi, R., Reich, D., 2015. Genome-wide patterns of selection in 230 ancient Eurasians. Nature 528, 499–503. https://doi.org/10.1038/nature16152

McColl, H., Racimo, F., Vinner, L., Demeter, F., Gakuhari, T., Moreno-Mayar, J.V., Driem, G. van, Wilken, U.G., Seguin-Orlando, A., Castro, C. de la F., Wasef, S., Shoocongdej, R., Souksavatdy, V., Sayavongkhamdy, T., Saidin, M.M., Allentoft, M.E., Sato, T., Malaspinas, A.-S., Aghakhanian, F.A., Korneliussen, T., Prohaska, A., Margaryan, A., Damgaard, P. de B., Kaewsutthi, S., Lertrit, P., Nguyen, T.M.H., Hung, H., Tran, T.M., Truong, H.N., Nguyen, G.H., Shahidan, S., Wiradnyana, K., Matsumae, H., Shigehara, N., Yoneda, M., Ishida, H., Masuyama, T., Yamada, Y., Tajima, A., Shibata, H., Toyoda, A., Hanihara, T., Nakagome, S., Deviese, T., Bacon, A.-M., Duringer, P., Ponche, J.-L., Shackelford, L., Patole-Edoumba, E., Nguyen, A.T., Bellina-Pryce, B., Galipaud, J.-C., Kinaston, R., Buckley, H., Pottier, C., Rasmussen, S., Higham, T., Foley, R.A., Lahr, M.M., Orlando, L., Sikora, M., Phipps, M.E., Oota, H., Higham, C., Lambert, D.M., Willerslev, E., 2018. The prehistoric peopling of Southeast Asia. Science 361, 88–92. https://doi.org/10.1126/science.aat3628

Narasimhan, V.M., Patterson, N., Moorjani, P., Rohland, N., Bernardos, R., Mallick, S., Lazaridis, I., Nakatsuka, N., Olalde, I., Lipson, M., Kim, A.M., Olivieri, L.M., Coppa, A., Vidale, M., Mallory, J., Moiseyev, V., Kitov, E., Monge, J., Adamski, N., Alex, N., Broomandkhoshbacht, N., Candilio, F., Callan, K., Cheronet, O., Culleton, B.J., Ferry, M., Fernandes, D., Freilich, S., Gamarra, B., Gaudio, D., Hajdinjak, M., Harney, É., Harper, T.K., Keating, D., Lawson, A.M., Mah, M., Mandl, K., Michel, M., Novak, M., Oppenheimer, J., Rai, N., Sirak, K., Slon, V., Stewardson, K., Zalzala, F., Zhang, Z., Akhatov, G., Bagashev, A.N., Bagnera, A., Baitanayev, B., Bendezu-Sarmiento, J., Bissembaev, A.A., Bonora, G.L., Chargynov, T.T., Chikisheva, T., Dashkovskiy, P.K., Derevianko, A., Dobeš, M., Douka, K., Dubova, N., Duisengali, M.N., Enshin, D., Epimakhov, A., Fribus, A.V., Fuller, D., Goryachev, A., Gromov, A., Grushin, S.P., Hanks, B., Judd, M., Kazizov, E., Khokhlov, A., Krygin, A.P., Kupriyanova, E., Kuznetsov, P., Luiselli, D., Maksudov, F., Mamedov, A.M., Mamirov, T.B., Meiklejohn, C., Merrett, D.C., Micheli, R., Mochalov, O., Mustafokulov, S., Nayak, A., Pettener, D., Potts, R., Razhev, D., Rykun, M., Sarno, S., Savenkova, T.M., Sikhymbaeva, K., Slepchenko, S.M., Soltobaev, O.A., Stepanova, N., Svyatko, S., Tabaldiev, K., Teschler-Nicola, M., Tishkin, A.A., Tkachev, V.V., Vasilyev, S., Velemínský, P., Voyakin, D., Yermolayeva, A., Zahir, M., Zubkov, V.S., Zubova, A., Shinde, V.S., Lalueza-Fox, C., Meyer, M., Anthony, D., Boivin, N., Thangaraj, K., Kennett, D.J., Frachetti, M., Pinhasi, R., Reich, D., 2019. The formation of human populations in South and Central Asia. Science 365, eaat7487. https://doi.org/10.1126/science.aat7487

Olalde, I., Brace, S., Allentoft, M.E., Armit, I., Kristiansen, K., Booth, T., Rohland, N., Mallick, S., Szécsényi-Nagy, A., Mittnik, A., Altena, E., Lipson, M., Lazaridis, I., Harper, T.K., Patterson, N., Broomandkhoshbacht, N., Diekmann, Y., Faltyskova, Z., Fernandes, D., Ferry, M., Harney, E., de Knijff, P., Michel, M., Oppenheimer, J., Stewardson, K., Barclay, A., Alt, K.W., Liesau, C., Ríos, P., Blasco, C., Miguel, J.V., García, R.M., Fernández, A.A., Bánffy, E., Bernabò-Brea, M., Billoin, D., Bonsall, C., Bonsall, L., Allen, T., Büster, L., Carver, S., Navarro, L.C., Craig, O.E., Cook, G.T., Cunliffe, B., Denaire, A., Dinwiddy, K.E., Dodwell, N., Ernée, M., Evans, C., Kuchařík, M., Farré, J.F., Fowler, C., Gazenbeek, M., Pena, R.G., Haber-Uriarte, M., Haduch, E., Hey, G., Jowett, N., Knowles, T., Massy, K., Pfrengle, S., Lefranc, P., Lemercier, O., Lefebvre, A., Martínez, C.H., Olmo, V.G., Ramírez, A.B., Maurandi, J.L., Majó, T., McKinley, J.I., McSweeney, K., Mende, B.G., Mod, A., Kulcsár, G., Kiss, V., Czene, A., Patay, R., Endrődi, A., Köhler, K., Hajdu, T., Szeniczey, T., Dani, J., Bernert, Z., Hoole, M., Cheronet, O., Keating, D., Velemínský, P., Dobeš, M., Candilio, F., Brown, F., Fernández, R.F., Herrero-Corral, A.-M., Tusa, S., Carnieri, E., Lentini, L., Valenti, A., Zanini, A., Waddington, C., Delibes, G., Guerra-Doce, E., Neil, B., Brittain, M., Luke, M., Mortimer, R., Desideri, J., Besse, M., Brücken, G., Furmanek, M., Hałuszko, A., Mackiewicz, M., Rapiński, A., Leach, S., Soriano, I., Lillios, K.T., Cardoso, J.L., Pearson, M.P., Włodarczak, P., Price, T.D., Prieto, P., Rey, P.-J., Risch, R., Rojo Guerra, M.A., Schmitt, A., Serralongue, J., Silva, A.M., Smrčka, V., Vergnaud, L., Zilhão, J., Caramelli, D., Higham, T., Thomas, M.G., Kennett, D.J., Fokkens, H., Heyd, V., Sheridan, A., Sjögren, K.-G., Stockhammer, P.W., Krause, J., Pinhasi, R., Haak, W., Barnes, I., Lalueza-Fox, C., Reich, D., 2018. The Beaker phenomenon and the genomic transformation of northwest Europe. Nature 555, 190–196. https://doi.org/10.1038/nature25738

Olalde, I., Mallick, S., Patterson, N., Rohland, N., Villalba-Mouco, V., Silva, M., Dulias, K., Edwards, C.J., Gandini, F., Pala, M., Soares, P., Ferrando-Bernal, M., Adamski, N., Broomandkhoshbacht, N., Cheronet, O., Culleton, B.J., Fernandes, D., Lawson, A.M., Mah, M., Oppenheimer, J., Stewardson, K., Zhang, Z., Jiménez Arenas, J.M., Toro Moyano, I.J., Salazar-García, D.C., Castanyer, P., Santos, M., Tremoleda, J., Lozano, M., García Borja, P., Fernández-Eraso, J., Mujika-Alustiza, J.A., Barroso, C., Bermúdez, F.J., Viguera Mínguez, E., Burch, J., Coromina, N., Vivó, D., Cebrià, A., Fullola, J.M., García-Puchol, O., Morales, J.I., Oms, F.X., Majó, T., Vergès, J.M., Díaz-Carvajal, A., Ollich-Castanyer, I., López-Cachero, F.J., Silva, A.M., Alonso-Fernández, C., Delibes de Castro, G., Jiménez Echevarría, J., Moreno-Márquez, A., Pascual Berlanga, G., Ramos-García, P., Ramos-Muñoz, J., Vijande Vila, E., Aguilella Arzo, G., Esparza Arroyo, Á., Lillios, K.T., Mack, J., Velasco-Vázquez, J., Waterman, A., Benítez de Lugo Enrich, L., Benito Sánchez, M., Agustí, B., Codina, F., de Prado, G., Estalrrich, A., Fernández Flores, Á., Finlayson, C., Finlayson, G., Finlayson, S., Giles-Guzmán, F., Rosas, A., Barciela González, V., García Atiénzar, G., Hernández Pérez, M.S., Llanos, A., Carrión Marco, Y., Collado Beneyto, I., López-Serrano, D., Sanz Tormo, M., Valera, A.C., Blasco, C., Liesau, C., Ríos, P., Daura, J., de Pedro Michó, M.J., Diez-Castillo, A.A., Flores Fernández, R., Francès Farré, J., Garrido-Pena, R., Gonçalves, V.S., Guerra-Doce, E., Herrero-Corral, A.M., Juan-Cabanilles, J., López-Reyes, D., McClure, S.B., Merino Pérez, M., Oliver Foix, A., Sanz Borràs, M., Sousa, A.C., Vidal Encinas, J.M., Kennett, D.J., Richards, M.B., Werner Alt, K., Haak, W., Pinhasi, R., Lalueza-Fox, C., Reich, D., 2019. The genomic history of the Iberian Peninsula over the past 8000 years. Science 363, 1230–1234. https://doi.org/10.1126/science.aav4040

Posth, C., Nakatsuka, N., Lazaridis, I., Skoglund, P., Mallick, S., Lamnidis, T.C., Rohland, N., Nägele, K., Adamski, N., Bertolini, E., Broomandkhoshbacht, N., Cooper, A., Culleton, B.J., Ferraz, T., Ferry, M., Furtwängler, A., Haak, W., Harkins, K., Harper, T.K., Hünemeier, T., Lawson, A.M., Llamas, B., Michel, M., Nelson, E., Oppenheimer, J., Patterson, N., Schiffels, S., Sedig, J., Stewardson, K., Talamo, S., Wang, C.-C., Hublin, J.-J., Hubbe, M., Harvati, K., Nuevo Delaunay, A., Beier, J., Francken, M., Kaulicke, P., Reyes-Centeno, H., Rademaker, K., Trask, W.R., Robinson, M., Gutierrez, S.M., Prufer, K.M., Salazar-García, D.C., Chim, E.N., Müller Plumm Gomes, L., Alves, M.L., Liryo, A., Inglez, M., Oliveira, R.E., Bernardo, D.V., Barioni, A., Wesolowski, V., Scheifler, N.A., Rivera, M.A., Plens, C.R., Messineo, P.G., Figuti, L., Corach, D., Scabuzzo, C., Eggers, S., DeBlasis, P., Reindel, M., Méndez, C., Politis, G., Tomasto-Cagigao, E., Kennett, D.J., Strauss, A., Fehren-Schmitz, L., Krause, J., Reich, D., 2018. Reconstructing the Deep Population History of Central and South America. Cell 175, 1185-1197.e22. https://doi.org/10.1016/j.cell.2018.10.027

Saag, Lehti, Laneman, M., Varul, L., Malve, M., Valk, H., Razzak, M.A., Shirobokov, I.G., Khartanovich, V.I., Mikhaylova, E.R., Kushniarevich, A., Scheib, C.L., Solnik, A., Reisberg, T., Parik, J., Saag, Lauri, Metspalu, E., Rootsi, S., Montinaro, F., Remm, M., Mägi, R., D’Atanasio, E., Crema, E.R., Díez-del-Molino, D., Thomas, M.G., Kriiska, A., Kivisild, T., Villems, R., Lang, V., Metspalu, M., Tambets, K., 2019. The Arrival of Siberian Ancestry Connecting the Eastern Baltic to Uralic Speakers further East. Current Biology S0960982219304245. https://doi.org/10.1016/j.cub.2019.04.026

Sánchez-Quinto, F., Malmström, H., Fraser, M., Girdland-Flink, L., Svensson, E.M., Simões, L.G., George, R., Hollfelder, N., Burenhult, G., Noble, G., Britton, K., Talamo, S., Curtis, N., Brzobohata, H., Sumberova, R., Götherström, A., Storå, J., Jakobsson, M., 2019. Megalithic tombs in western and northern Neolithic Europe were linked to a kindred society. PNAS 116, 9469–9474. https://doi.org/10.1073/pnas.1818037116

Scheib, C.L., Li, H., Desai, T., Link, V., Kendall, C., Dewar, G., Griffith, P.W., Mörseburg, A., Johnson, J.R., Potter, A., Kerr, S.L., Endicott, P., Lindo, J., Haber, M., Xue, Y., Tyler-Smith, C., Sandhu, M.S., Lorenz, J.G., Randall, T.D., Faltyskova, Z., Pagani, L., Danecek, P., O’Connell, T.C., Martz, P., Boraas, A.S., Byrd, B.F., Leventhal, A., Cambra, R., Williamson, R., Lesage, L., Holguin, B., Ygnacio-De Soto, E., Rosas, J., Metspalu, M., Stock, J.T., Manica, A., Scally, A., Wegmann, D., Malhi, R.S., Kivisild, T., 2018. Ancient human parallel lineages within North America contributed to a coastal expansion. Science 360, 1024–1027. https://doi.org/10.1126/science.aar6851

Schroeder, H., Margaryan, A., Szmyt, M., Theulot, B., Włodarczak, P., Rasmussen, S., Gopalakrishnan, S., Szczepanek, A., Konopka, T., Jensen, T.Z.T., Witkowska, B., Wilk, S., Przybyła, M.M., Pospieszny, Ł., Sjögren, K.-G., Belka, Z., Olsen, J., Kristiansen, K., Willerslev, E., Frei, K.M., Sikora, M., Johannsen, N.N., Allentoft, M.E., 2019. Unraveling ancestry, kinship, and violence in a Late Neolithic mass grave. PNAS 116, 10705–10710. https://doi.org/10.1073/pnas.1820210116

Sikora, M., Pitulko, V.V., Sousa, V.C., Allentoft, M.E., Vinner, L., Rasmussen, S., Margaryan, A., Damgaard, P. de B., Fuente, C. de la, Renaud, G., Yang, M.A., Fu, Q., Dupanloup, I., Giampoudakis, K., Nogués-Bravo, D., Rahbek, C., Kroonen, G., Peyrot, M., McColl, H., Vasilyev, S.V., Veselovskaya, E., Gerasimova, M., Pavlova, E.Y., Chasnyk, V.G., Nikolskiy, P.A., Gromov, A.V., Khartanovich, V.I., Moiseyev, V., Grebenyuk, P.S., Fedorchenko, A.Y., Lebedintsev, A.I., Slobodin, S.B., Malyarchuk, B.A., Martiniano, R., Meldgaard, M., Arppe, L., Palo, J.U., Sundell, T., Mannermaa, K., Putkonen, M., Alexandersen, V., Primeau, C., Baimukhanov, N., Malhi, R.S., Sjögren, K.-G., Kristiansen, K., Wessman, A., Sajantila, A., Lahr, M.M., Durbin, R., Nielsen, R., Meltzer, D.J., Excoffier, L., Willerslev, E., 2019. The population history of northeastern Siberia since the Pleistocene. Nature 570, 182. https://doi.org/10.1038/s41586-019-1279-z

Skourtanioti, E., Erdal, Y.S., Frangipane, M., Balossi Restelli, F., Yener, K.A., Pinnock, F., Matthiae, P., Özbal, R., Schoop, U.-D., Guliyev, F., Akhundov, T., Lyonnet, B., Hammer, E.L., Nugent, S.E., Burri, M., Neumann, G.U., Penske, S., Ingman, T., Akar, M., Shafiq, R., Palumbi, G., Eisenmann, S., D’Andrea, M., Rohrlach, A.B., Warinner, C., Jeong, C., Stockhammer, P.W., Haak, W., Krause, J., 2020. Genomic History of Neolithic to Bronze Age Anatolia, Northern Levant, and Southern Caucasus. Cell 181, 1158-1175.e28. https://doi.org/10.1016/j.cell.2020.04.044

Wang, C.-C., Reinhold, S., Kalmykov, A., Wissgott, A., Brandt, G., Jeong, C., Cheronet, O., Ferry, M., Harney, E., Keating, D., Mallick, S., Rohland, N., Stewardson, K., Kantorovich, A.R., Maslov, V.E., Petrenko, V.G., Erlikh, V.R., Atabiev, B.Ch., Magomedov, R.G., Kohl, P.L., Alt, K.W., Pichler, S.L., Gerling, C., Meller, H., Vardanyan, B., Yeganyan, L., Rezepkin, A.D., Mariaschk, D., Berezina, N., Gresky, J., Fuchs, K., Knipper, C., Schiffels, S., Balanovska, E., Balanovsky, O., Mathieson, I., Higham, T., Berezin, Y.B., Buzhilova, A., Trifonov, V., Pinhasi, R., Belinskij, A.B., Reich, D., Hansen, S., Krause, J., Haak, W., 2019. Ancient human genome-wide data from a 3000-year interval in the Caucasus corresponds with eco-geographic regions. Nat Commun 10, 590. https://doi.org/10.1038/s41467-018-08220-8
